# Kinetic asymmetry drives kinesin-1’s unidirectional and processive movement

**DOI:** 10.1101/2025.06.11.659012

**Authors:** Hiroshi Isojima, Kohei Matsuzaki, Michio Tomishige

## Abstract

Kinesin-1 is a dimeric motor protein that uses ATP hydrolysis energy to move along microtubules in a hand-over-hand manner^1^. The unidirectional movement of kinesin-1 has traditionally been explained by an ATP-dependent power stroke action of the neck linker^2–4^, connecting its two catalytic domains (heads), that biases the diffusional motion forward (biased-diffusion). However, recent studies on synthetic molecular motors have supported a Brownian ratchet mechanism based on kinetic asymmetry between two locations (biased- binding)^5–9^, and which of these mechanisms applies to biological motors remains debated^10–12^. Here, we engineered a two-headed kinesin that alternately uses these mechanisms to step forward, allowing us to investigate how they contribute to unidirectional movement. The tethered head that uses biased-diffusion frequently rebound to the rear-binding site but eventually stepped forward, as the front head remained securely bound to the microtubule. The biased-binding mechanism proved more efficient by preventing rebinding of the detached head and was independent of ATP binding. Instead, ATP hydrolysis energy is primarily consumed to ensure preferential detachment of the rear head. These findings demonstrate that kinesin-1 functions as an information ratchet based on kinetic asymmetry in microtubule-binding and detachment of the heads, while power strokes serve to enhance movements under load.

---

Biological molecular motors convert chemical energy into unidirectional movement, despite operating in an environment dominated by Brownian motion^13^. Kinesin-1 (hereafter referred to as kinesin) is a dimeric motor protein that transports cargo inside cells by moving toward the plus-end of the microtubule^1^. Kinesin moves processively for about one micrometer before dissociating from the microtubule and can resist up to 7 pN hindering load^14^. The unidirectional motion of kinesin occurs through a hand-over-hand movement of two motor domains (“heads”), powered by ATP hydrolysis in the catalytic head^15–17^. ATP hydrolysis and subsequent Pi release cause the trailing head to detach from the microtubule. The detached head remains tethered and unbound until ATP binds to its partner head on the microtubule, which triggers the tethered head’s binding to the forward-tubulin binding site^18–22^.

The neck linker docking model^3,4^ is the most widely accepted explanation for ATP-induced stepping. According to this model, ATP binding causes the neck linker—a 14 amino-acid segment connecting the two heads—to dock onto the microtubule-bound head, creating a power stroke (energetically downhill conformational change) that moves the tethered head forward by about 4 nm (Fig. 1a, left). While this translocation alone is not sufficient for stepping, it works in concert with Brownian motion: the neck linker docking shifts the center of the diffusional movement of the tethered head forward^23^, promoting its attachment to the forward-tubulin binding site^4^. This mechanism, which we termed the “biased-diffusion” mechanism, represents a type of Brownian ratchet known as an “energy ratchet”^8,13^. Furthermore, after ATP binding, a 9-amino-acid N-terminal region called the “cover-strand” forms a parallel β-sheet with part of the neck linker. This interaction guides the tethered head forward before the neck linker docking is complete, particularly when walking against hindering load^24,25^. However, these mechanisms do not explain what prevents the tethered head from rebinding to the rear-tubulin binding site especially before ATP binding.

**Figure 1.**
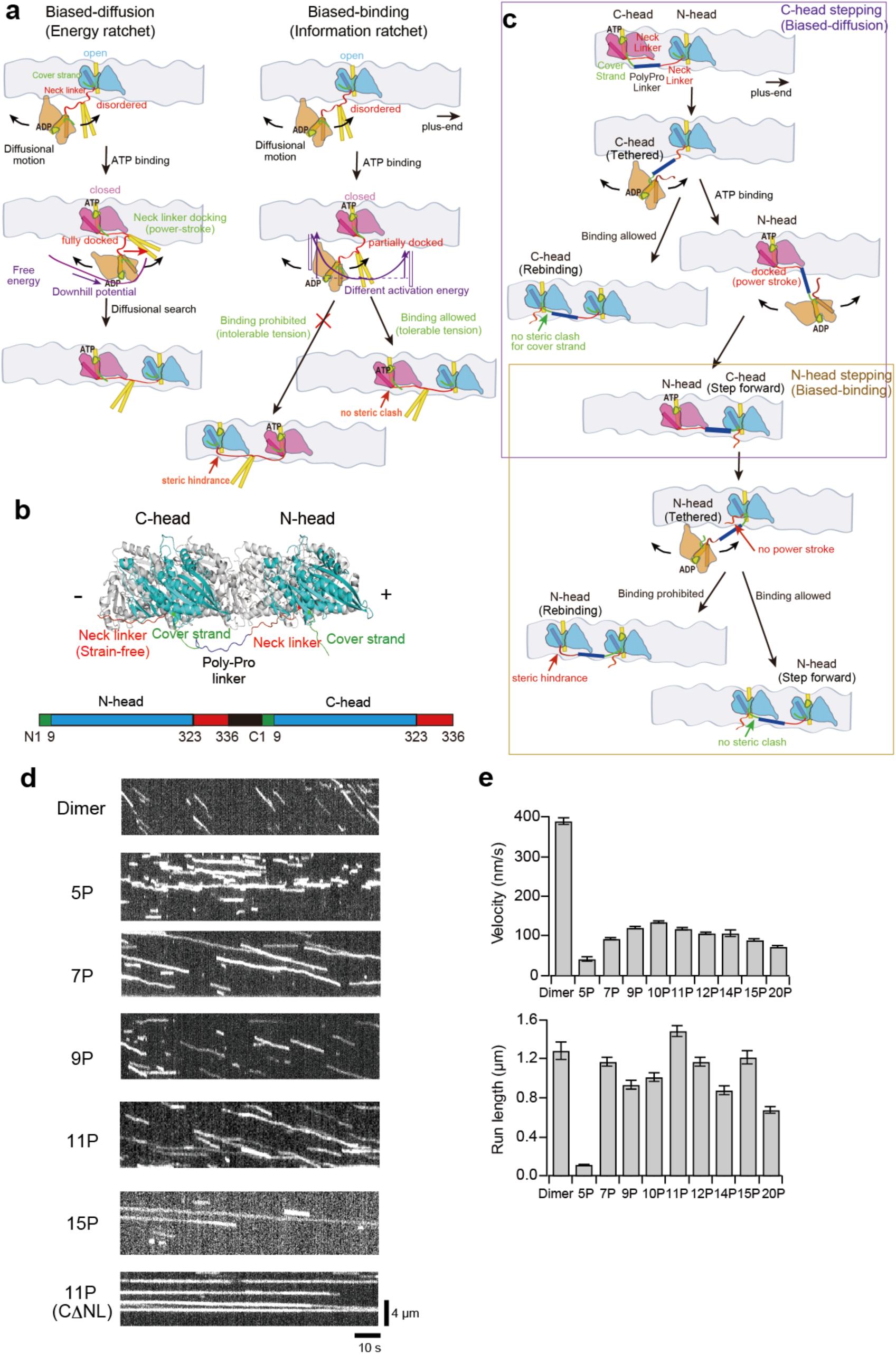
Motility cycle and single-molecule movement of dimeric and tandemly-joined two-headed (tandem) kinesin along microtubules. (**a**) Schematics showing the biased-diffusion and biased-binding mechanisms to explain the preferential forward stepping of dimeric kinesin. Three distinct conformational states of the head are shown in cyan (open state), magenta (closed state), and orange (semi-open state), with the α6 helix, which connects to the neck linker, highlighted as a rod. The cover strand and neck linker are depicted in green and red, while the α4 helix, which directly interacts with the microtubule, and the neck coiled-coil are shown in yellow. Purple lines indicate the free energy landscapes of the tethered head. (**b**) Atomic model of the tandem kinesin in the two-head-bound state (top view), where the N-terminal head (“N-head”) leads and the C-terminal head (“C-head”) trails (both in open state; side view is shown in Supplementary Fig. 1b). Since the neck linker of the C-head is not connected to the N-head, it remains free of strain. (**c**) Schematic showing the hand-over-hand movement of tandem kinesin, which alternates between biased-diffusion and biased-binding mechanisms to step forward. (**d**) Kymographs showing the motility of Cy5-labeled tandem kinesins with different linker length (number of prolines) along axonemes in the presence of 1 mM ATP. CΔNL denotes an 11P tandem kinesin whose C-head’s neck linker (residues 324-336) has been deleted. Dimeric kinesin (490 amino-acids) is shown for comparison. (**e**) Average velocity and run length (± s.e.m.) of tandem kinesin at 1 mM ATP. Histograms are shown in Supplementary Fig. 2b.

We recently proposed a neck-linker tension-based regulation mechanism to explain how the rebinding of the tethered is prohibited^26^. In this model, microtubule binding is asymmetrically regulated; the tethered head is prohibited from binding to the rear-binding site due to excessive neck linker tension, while it is allowed to bind to the forward-binding site after ATP binding. This asymmetry arises from steric hindrance on the surface of the trailing nucleotide-free head, a bulge just ahead of the neck linker’s base which interferes neck linker stretching specifically in the forward direction^26^. The difference in the neck linker tension (and thus entropy reduction of the stretched neck linkers) creates a kinetic asymmetry between forward and rearward microtubule binding sites (Fig. 1a, right). This “biased-binding” mechanism represents another type of Brownian ratchet called an “information ratchet”^8,13^. The unbound head’s behavior depends on information about its current state, specifically whether the head occupies a forward or rearward binding site. This position affects the neck linker tension, which in turn modulates the activation energy and microtubule-binding rate.

Since the biased-diffusion and biased-binding mechanisms operate in distinct phases, they are not mutually exclusive and likely both contribute to the preferential forward stepping. While distinguishing between them and measuring their relative contributions in native dimeric kinesin remains challenging, synthetic molecular motors have provided simpler model systems for understanding motor function under thermal fluctuations. Recent studies with chemically driven synthetic motors demonstrated that kinetic preference for one direction over the other during the mechanochemical cycle (called “kinetic asymmetry”) is essential for directionality, while power strokes are not essential but rather enhance movement efficiency^27–29^. Whether these principles apply to more complex biological molecular motors remains a matter of debate.

## Design of tandemly joined two-headed kinesin

To differentiate between the biased-diffusion and biased-binding mechanisms, we focused on the distinct roles of the neck linkers in the tethered and microtubule-bound heads. The biased diffusion mechanism requires that the neck linker of the “microtubule-bound head” connects to the tethered head, allowing it to transmit the conformational change that biases forward diffusion (Fig. 1a, left). In contrast, the biased binding mechanism requires that the neck linker of the “tethered head” connects to the microtubule-bound head to detect steric hindrance (Fig. 1a, right). To investigate how each mechanism contributes to kinesin stepping, we engineered a two-headed monomer construct by joining two cysteine-light monomeric heads (336 amino-acids) in tandem (called “tandem kinesin”). In this construct, the neck linker of the N-terminal head (“N-head”) connects to the cover strand of the C-terminal head (“C-head”), while the neck linker of C-head is unconnected and free of strain (Fig. 1b and Supplementary Fig. 1a). Since the cover strand is shorter than the neck linker, we inserted rigid poly-proline spacers of varying lengths between the N-head’s neck linker and the C-head’s cover strand. If this tandem kinesin moves along the microtubule in a hand-over-hand manner, the tethered C-head advances solely through the biased-diffusion mechanism to step forward, as its neck linker is not connected to the microtubule-bound N-head, preventing it from detecting tension increases caused by the steric hindrance just ahead of the neck linker base (Fig. 1c and Supplementary Fig. 1b). Conversely, the tethered N-head relies exclusively on the biased-binding mechanism, as the microtubule-bound C-head’s neck linker is not connected to the N-head and thus cannot propel it through neck linker docking.

## Tandem kinesins exhibit processive movement

First, we investigated whether the tandem kinesin could move processively and unidirectionally along microtubules. Using total internal reflection microscopy, we observed fluorescently labeled tandem kinesin bound to the microtubule in the presence of 1 mM ATP (Fig. 1d and Supplementary Fig. 2). The tandem kinesin with 5 poly-Pro insertion (termed 5P) exhibited one-dimensional diffusional movement with a slight plus-end bias. Tandem kinesin with 7 or more poly-Pro insertions showed unidirectional movement. The velocity increased with insertion length up to 10P residues but decreased with lengths beyond 11P (Fig. 1e). The run length remained relatively constant across different insertion sizes, reaching its maximum for 11P (Fig. 1e). Our results, which demonstrate tandem kinesin’s unidirectional and processive movement, show that forward stepping can be achieved through either the biased-diffusion or biased-binding mechanism alone. When we deleted the neck linker from the C-head (from residue 324) of 11P tandem kinesin (termed CΔNL), the resulting molecule bound to the microtubule but remained stationary (Fig. 1d), consistent with the previous finding that the neck linker docking is essential for promoting ATP hydrolysis of the trailing head^30–32^.

Having demonstrated that the 11P construct achieved the maximum run length, we focused our subsequent investigations on the detailed motion of this 11P tandem kinesin. We first compared the motility and ATPase activity of 11P tandem kinesin with wild-type dimeric kinesin. The ATP turnover rate (40 s⁻¹ per head; Supplementary Fig. 3a, b) and the run length (1.49 µm; Fig. 1e) of 11P tandem kinesin were comparable to those of wild-type dimer (49 s⁻¹ per head and 1.29 µm, respectively). However, the velocity of 11P was more than three times slower than that of the dimer (117 and 387 nm/s for tandem and dimer, respectively). These findings suggest that the coupling between ATP hydrolysis and forward step was less efficient than in wild-type dimers (Supplementary Fig. 3c). This inefficiency resembles the behavior of neck linker extended mutants, which showed impaired head-head coordination^21,33,34^.

## Alternating fast and slow steps

To examine whether tandem kinesin moves in a hand-over-hand manner, we employed a single-molecule FRET sensor that could distinguish between the two two-head-bound states^18^. We introduced two cysteine residues into the cysteine-light 11P tandem kinesin; one at the minus-end oriented base of the N-head (43Cys) and another at the plus-end oriented tip of the C-head (215Cys) (Supplementary Fig. 4a). These cysteines were labeled with Cy3 (donor) and Cy5 (acceptor) fluorophores. When both heads bind to adjacent tubulin-binding sites along a single protofilament, we expect to observe a high FRET efficiency (∼90%) when the N-head leads, and a low FRET efficiency (∼10%) when the C-head leads (Fig. 2a and Supplementary Fig. 4b). In the one-head-bound state, we expect an intermediate FRET efficiency (∼50%; Supplementary Fig. 4b). In the presence of 1 mM ATP, the dual-labeled molecules showed two-state transition between high and low FRET states (Fig. 2b), with transitions towards the low FRET state occurring approximately once per 16 nm step (Fig. 2c). These results demonstrate that, like dimeric kinesin, the 11P tandem kinesin moves in a hand-over-hand manner.

**Figure 2.**
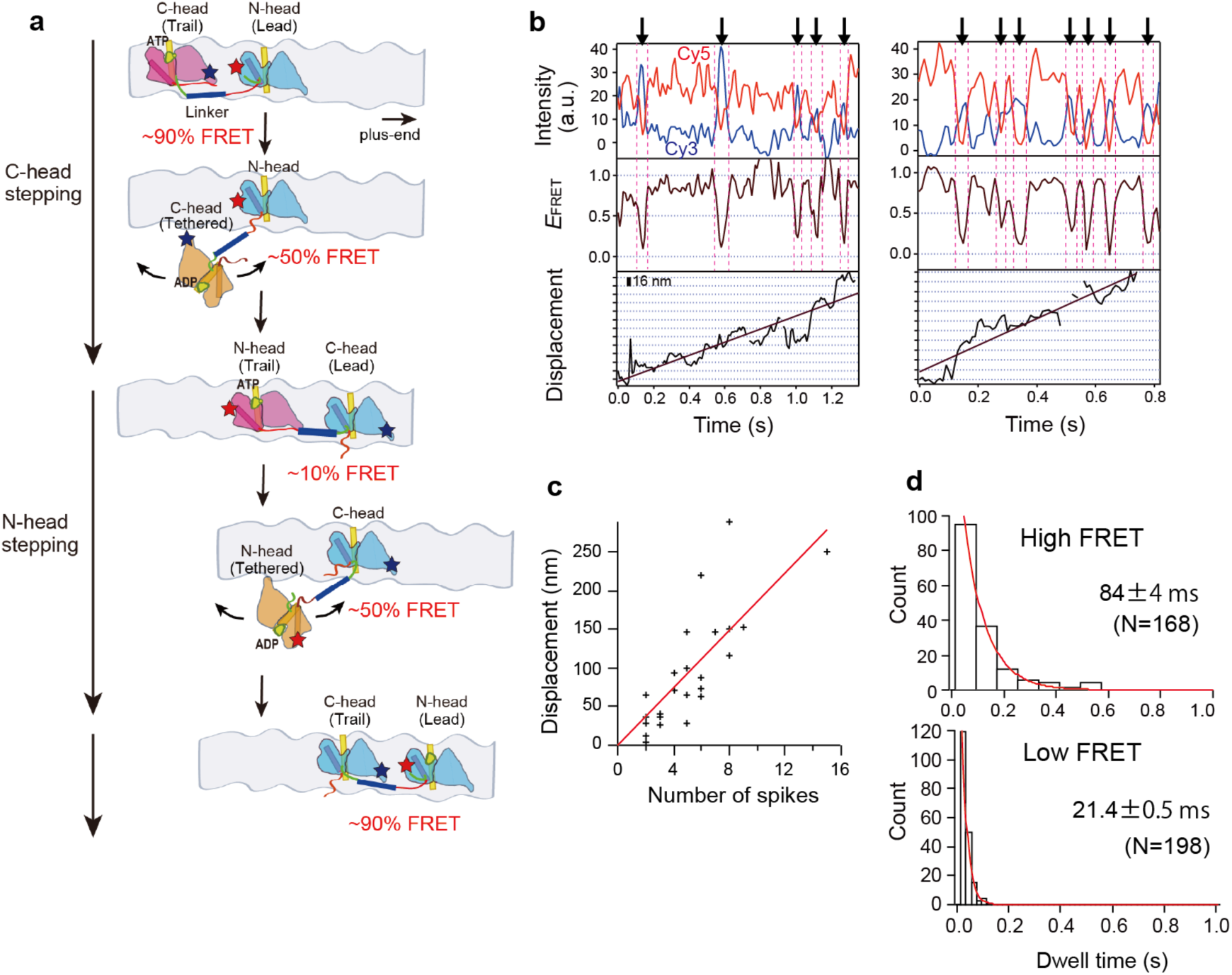
SmFRET observations of head-head configuration of tandem kinesin during processive motility. (**a**) Diagram illustrating FRET efficiency changes during the hand-over-hand movement of tandem kinesin, with fluorescent dyes attached to the rear of N-head (C43; Supplementary Fig. 4a) and front of C-head (C215). High (∼90%) FRET efficiency indicates the N-head is leading in the two-head-bound state, while low (∼10%) FRET efficiency indicates the C-head is leading. (**b**) Representative traces of fluorescence intensities of donor (Cy3, blue) and acceptor (Cy5, red) fluorophores, FRET efficiency and axial displacement for Cy3/Cy5-labeled tandem kinesin at 1 mM ATP. Arrows indicate transitions to low FRET states. (**c**) Relationship between number of transitions towards the low FRET state and the displacement per observation time; each point represents a different molecule. The red line shows a linear fit (18.3 nm per low FRET transition). (**d**) Distributions of dwell times in the high FRET and low FRET states. Red lines show exponential fit. Numbers show mean dwell time (± s.e.m.) obtained by the fit. Numbers in parentheses indicate the number of dwells analyzed.

However, the FRET efficiency traces revealed alternating dwell times between steps, with the high FRET state lasting approximately four times longer than the low FRET state under saturating ATP conditions (Fig. 2d). We confirmed this finding using another smFRET sensor with reversed labeling positions (Supplementary Fig. 5). This difference became even more pronounced under limited ATP conditions (Supplementary Fig. 6 and 7). Since the dwell time of the high FRET state corresponds to the time required for the C-head to detach from the rear-binding site and binds to the forward-binding site (Fig. 2a), these findings indicate that the rear C-head steps forward less efficiently than the N-head.

## Detached C-head frequently rebound

The slower stepping of the C-head could be attributed to three factors: slow detachment of the rear C-head from the microtubule, frequent rebinding of the detached C-head to the rear tubulin-binding site, or extended dwell time of the detached C-head. To investigate these possibilities, we observed microtubule-binding and detachment of individual heads using total internal reflection dark-field microscopy with 50 µs temporal resolution^21^. We attached a 40-nm diameter gold nanoparticle to one head (via 55Cys) and observed the gold-labeled 11P tandem kinesin on the microtubule in the presence of 1 mM ATP (Fig. 3a). Gold probes attached to either the N-head or C-head showed stepwise movements of approximately 17 nm (Fig. 3b, c), while probes attached to the poly-Pro linker showed 8.3 nm steps (Fig. 3c and Supplementary Fig. 8). These findings further support that tandem kinesin moves in a hand-over-hand manner, where the center of mass steps match the spacing between adjacent tubulin-binding sites (8.3 nm), while steps of individual head are twice this distance.

**Figure 3.**
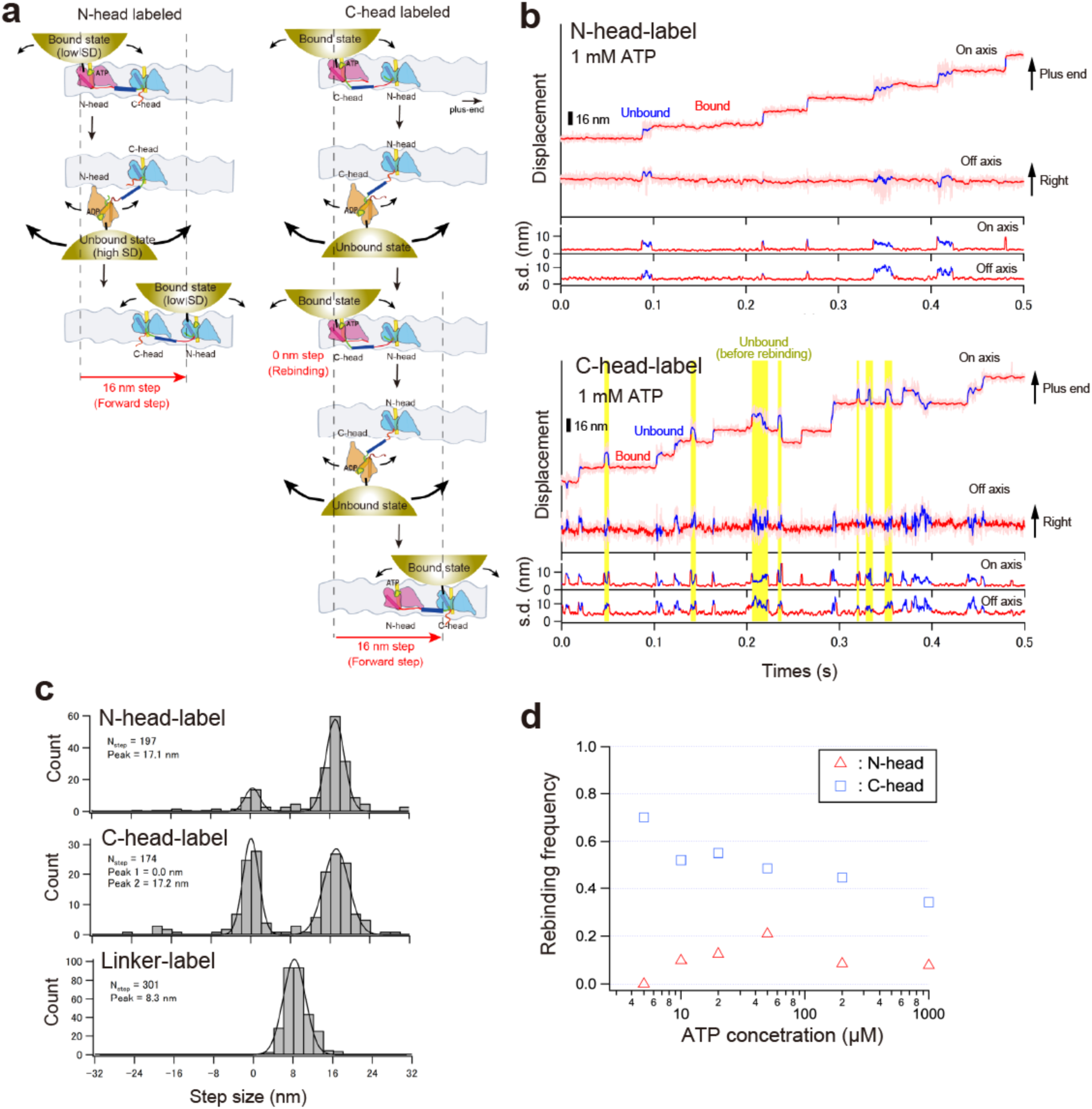
High-speed single-molecule observations of the gold probe attached to N- or C- head of tandem kinesin during processive motility. (**a**) Diagram showing the displacements and fluctuations of the gold probe attached to N- and C-head of tandem kinesin moving along microtubules. (**b**) Typical traces for the centroid positions of the gold probe attached to N- and C-head of 11P tandem kinesin (light red lines), toward the microtubule long axis (on axis) and perpendicular to the microtubule axis (off axis) in the presence of 1 mM ATP (recorded at 20,000 fps). Red and blue lines depict the median-filtered traces (window size of 51 frames) for the bound and unbound states, respectively. Lower panel shows the standard deviation (s.d.) of on- and off-axis positions for each time frame t (calculated as [t - 20, t + 20]). Yellow rectangles indicate the unbound states that are followed by rebinding to the rear-binding site (“rebinding” in panel **a**). (**c**) Histograms of the step sizes of gold probe attached to N-head, C-head, and poly- Pro linker (Supplementary Fig. 8) in the presence of 1 mM ATP. The average step sizes determined by fitting with Gaussian distributions (solid line) are shown. (**d**) The rebinding ratio calculated by dividing the number of observed rebinding events by the total number of rear head detachment events (see Supplementary Fig. 11 for detailed event counts).

This measurement also enables us to detect when the detached head rebinds to its original binding site (resulting in a 0 nm step (highlighted in yellow in Fig. 3b), because the 40-nm gold probe attached to the unbound head amplifies its diffusional movement through a rocking motion, allowing us to identify the head’s unbound states from the transient increases in the fluctuation^21,30^ (Fig. 3a)—something not possible with smaller probes. While the N-head-labeled tandem kinesin predominantly made 17-nm forward steps, the C-head-labeled tandem kinesin frequently exhibited 0-nm steps (Fig. 3c), indicating that the detached C-head often rebinds to its original rear binding site. This rebinding behavior explains both the C-head’s slower stepping rate and the tandem kinesin’s futile ATP hydrolysis. The frequency of rebinding of the detached rear N-head remained low (7.7% on average) and nearly independent of ATP concentration (Fig. 3d and Supplementary Figs. 9-11), while the detached rear C-head exhibited a higher rebinding frequency at saturating ATP concentration (34%) and showed further increases as the ATP concentration was decreased (70% at 5 µM ATP; Fig. 3d). These findings demonstrate that the biased-binding mechanism effectively suppresses rebinding of the detached head to the rear-binding site, regardless of ATP binding.

## ATP binding is not required to step forward

Next, we analyzed how fast the detached N- and C-heads step forward (Fig. 4a). The dwell times of the unbound (highly fluctuating) state of both N- and C-heads were independent of ATP concentration (3.4 and 2.4 ms, respectively; Fig. 4b and Supplementary Fig. 12). The C-head frequently rebound to the rear-binding site, and its dwell time in the unbound state before rebinding was also ATP-independent and slightly longer (3.3 ms; Fig. 4b) than before forward stepping. The dwell time of 2-3 ms aligns with that of dimeric kinesin at saturating ATP concentration when ATP binding can occur spontaneously^21,35,36^. However, these times are significantly longer that the calculated first passage time (∼5 µs) for the tethered head to reach the forward-binding site by Brownian diffusion^33^. These observations suggest that the detached head is initially prevented from microtubule binding, regardless of the conformational state of the microtubule-bound head, and an allosteric conformational transition of the detached head must occur before microtubule binding is possible^37,38^ (Fig. 6). Moreover, these findings suggest that, unlike dimeric kinesin which requires ATP binding to step forward, ATP binding to the partner head is not required for the tethered N- and C-heads to bind to forward- and rearward-binding sites, respectively. This finding aligns with the tension-based regulation mechanism^26^, where tandem kinesin can adopt two-head-bound states where both heads are in the nucleotide-free state, specifically when the N-head leads and the C-head trails (Fig. 1c and Supplementary Figs. 1b and 4b).

**Figure 4.**
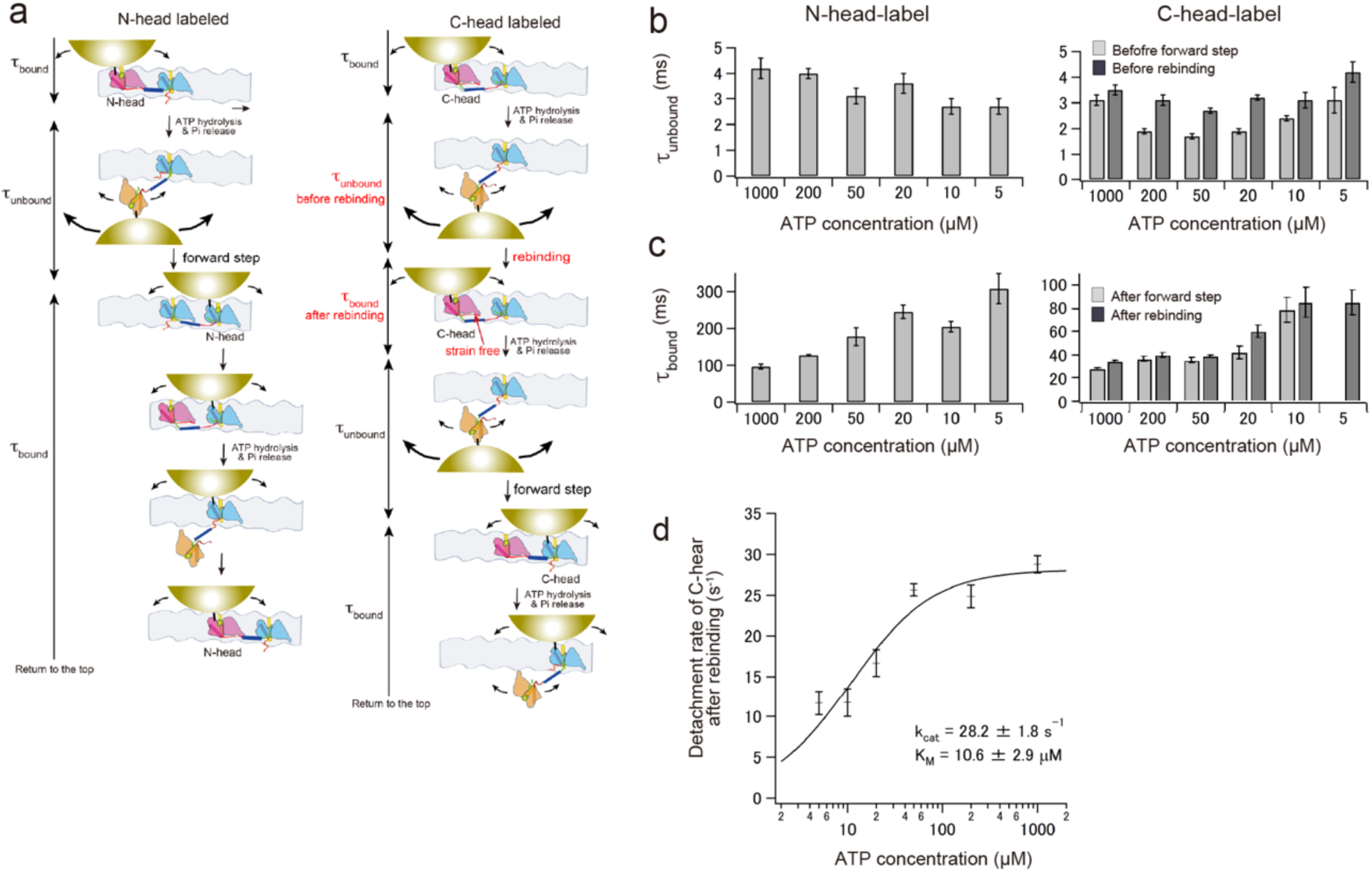
Dwell times of the microtubule unbound and bound states of N- or C-head of tandem kinesin during processive motility. (**a**) Diagram showing the dwell times in unbound (high s.d.) state and bound (low s.d.) state of gold-labeled N- and C-heads of tandem kinesin during processive motility. (**b**) Mean dwell times in the unbound state (1_unbound_) of N- and C- heads (histograms are shown in Supplementary Fig. 12). The 1_unbound_ of C-head was classified into periods before forward step and before rebinding (see panel **a**). Error bars represent s.e.m. (**c**) Mean dwell times in the bound state (1_bound_) of N- and C-heads (histograms are shown in Supplementary Fig. 14). The 1_bound_ of C-head was classified into periods after forward step and after rebinding. The dwell time histograms of C-head after the forward step were fitted with a double exponential function; one rate constant τ_2_ was fixed at 10.5 ms (corresponding to the dwell time of trailing N-head). The other rate constant τ_1_ (corresponding to trailing C-head) is shown here and is comparable to that of C-head after rebinding, which includes only the dwell time of trailing C-head. (**d**) Inverse of the average dwell time in the bound state 1_bound_ after rebinding were plotted as a function of ATP concentration. Error bars represent s.e.m. The solid line show the fit with the Michaelis-Menten equation and the fit parameters are shown.

## Slower detachment of trailing C-head

The tandem kinesin also provides insights into the gating mechanism in the two-head-bound state. To move in a hand-over-hand manner, the trailing head must hydrolyze ATP and detach from the microtubule before the leading head. Our recent kinetic studies^30^, along with cryo-EM studies by Sosa’s group^31,32^, demonstrated that backward strain on the neck linker of the leading head suppresses the allosteric conformational change from open (nucleotide-pocket opened) to closed (nucleotide-pocket closed, hydrolysis competent) conformational transition (front-head gating), while forward strain on the neck linker of the trailing head stabilizes the closed conformation and thereby promotes ATP hydrolysis and subsequent detachment from the microtubule (rear-head gating).

We first estimated the detachment rates of the trailing N- and C-heads from the microtubule to examine how neck linker strain affects these rates—the rear N-head experiences forward pulling force from the neck linker while the rear C-head does not (Supplementary Figs. 1b and 8a). To estimate the detachment rate of the trailing N-head, we employed the dwell times of the linker-labeled tandem kinesin. The gold probe attached to the poly-Pro linker showed alternating fast and slow steps (Supplementary Figs. 8b and 13a), consistent with the observations using smFRET (Fig. 2d and Supplementary Fig. 7b); short dwell time (low FRET state; Fig. 2a) represents the N-head is in the trailing position (Supplementary Fig. 8a). The dwell time before the fast step was independent of ATP concentration (Supplementary Fig. 13b, c), indicating that ATP must already be bound to the N-head by the time the partner C-head binds to the forward binding site, which aligns with our earlier observations of ATP-dependent C-head rebinding rates (Fig. 3d). The dwell time before the fast step averaged 10.9 ms, matching the previously observed dwell time (∼10 ms) of the trailing head in the wild-type dimer^21^. This demonstrates that the trailing N-head detaches as efficiently as the trailing head of dimeric kinesin.

The C-head frequently rebound to the rear-binding site, and the dwell time of the bound state after rebinding was shorter than that after the forward step (Supplementary Fig. 14). This difference occurs because the dwell time after rebinding only includes the trailing position of the labeled C-head, while the dwell time after forward step includes both dwells in the leading and trailing positions (Fig. 4a). Therefore, we used the dwell time after rebinding to estimate the C-head’s dwell time in the trailing position. The dwell time of the rear C-head depended on ATP concentration (Fig. 4c), and the inverse of the dwell time fit well with the Michaelis-Menten equation (Fig. 4d), yielding a maximum detachment rate of 28 s^-1^. This detachment rate of the rear C-head is approximately three times slower than that of the rear N-head calculated above (92 s^-1^; inverse of 10.9 ms), demonstrating that forward strain on the neck linker accelerates the rear head detachment (rear-head gating). The reduced detachment rate stems from the destabilization of neck linker docking onto the C- head, which we confirmed using a smFRET sensor that distinguishes between the neck linker’s docked and undocked state^39^ (Fig 5a, b, right and Supplementary Fig. 15, right). These findings support the idea that forward strain on the neck linker is necessary to stabilize the closed, hydrolysis-competent conformation^30^.

**Figure 5.**
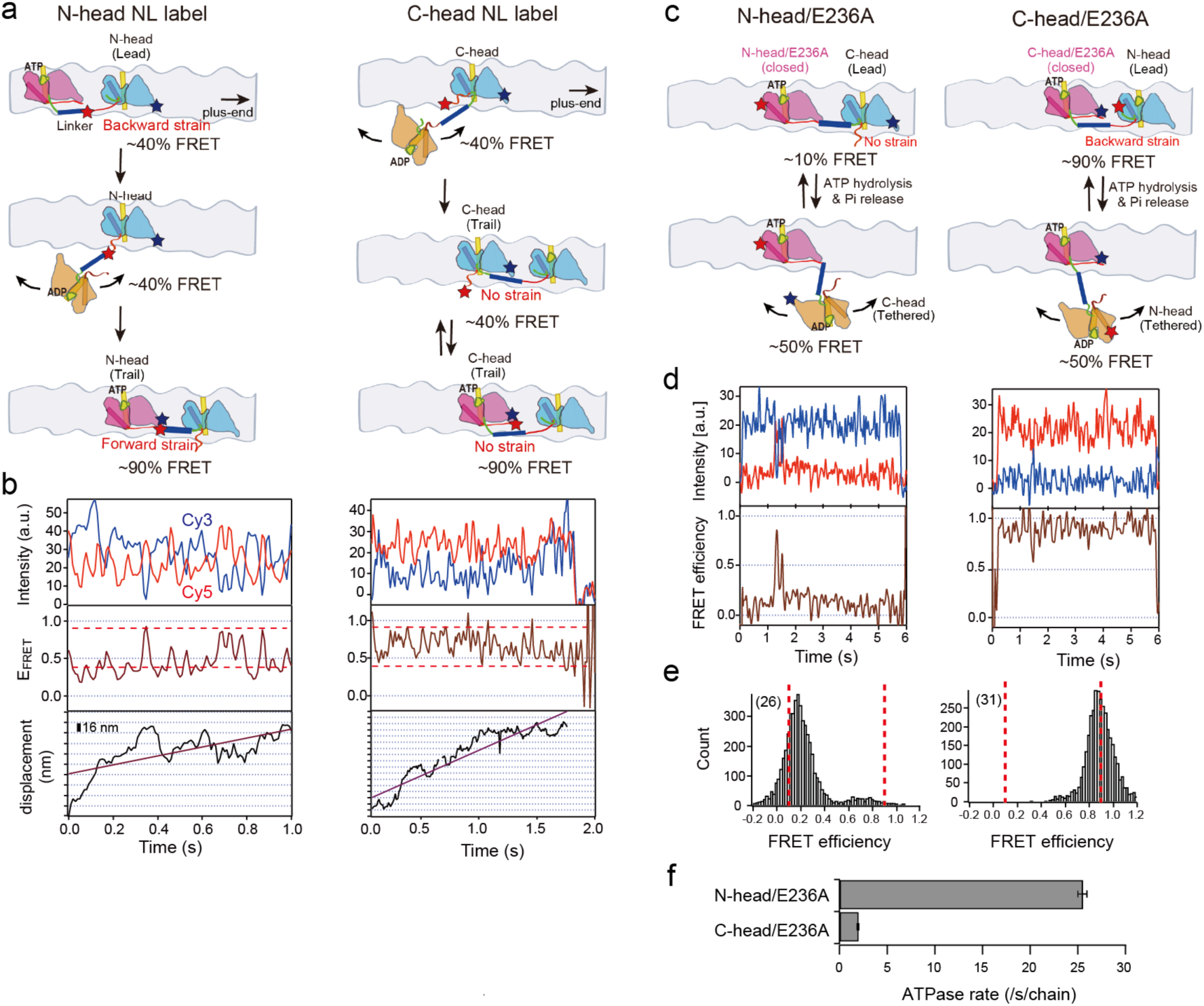
SmFRET observations of neck linker in tandem kinesin and head-head configurations in kinetically trapped tandem kinesin mutants. (**a**) Diagram illustrating the FRET sensor that is sensitive to neck linker (NL) conformational change attached to N- or C- head of tandem kinesin. The neck linker undocked and docked states can be distinguished as medium and high FRET states, respectively^39^ (positions for dye labeling are shown in Supplementary Fig. 15). (**b**) Typical traces of fluorescence intensities of donor (Cy3, blue) and acceptor (Cy5, red) fluorophores, FRET efficiency and axial displacement of Cy3/Cy5-labeled tandem kinesin at 1 mM ATP. Dotted lines illustrate neck linker docked and undocked states. During processive movement, the smFRET sensor on N-head (left) showed distinct two-state transitions similar to dimeric kinesin^39^. In contrast, the sensor on C-head displayed fluctuations between high and median FRET states (right), suggesting that its neck linker was not stable in either the closed or open state but rapidly switched between them. (**c**) Diagram illustrating microtubule detachment and binding of the leading head of tandem kinesin, where its partner head contains an E236A mutation that kinetically traps the head in a neck linker-docked closed state^3,30^. (**d**) Typical traces of fluorescence intensities of donor (Cy3, blue) and acceptor (Cy5, red) fluorophores, and FRET efficiency of Cy3/Cy5-labeled tandem kinesin at 1 mM ATP. (**e**) Histograms of the FRET efficiencies (from each frame of images) of dye-labeled tandem kinesin at 1 mM ATP. The numbers of molecules analyzed are shown in parentheses. Dotted lines illustrate peaks characteristic of two-head-bound states. (**f**) ATP turnover rates *k*_cat_. Since the catalytic activity of the E236 head was almost abolished^3^, the rates correspond to the activity of the partner head.

## Leading C-head is capable of ATP hydrolysis

When the C-head leads and N-head trails in the two-head-bound state of tandem kinesin, the neck linker of the leading C-head is not pulled backward and therefore is expected to be free of front-head gating (Fig. 1c). In our high-speed single-molecule observations of the C- head, we rarely observed detachment of the gold-labeled C-head when in the front position (11% of all detachment events (averaged across all ATP conditions); Supplementary Fig. 11). This occurs because the rear N-head, which is prebound by ATP and accelerated by the rear-head gating mechanism described above (detachment rate of ∼100 s⁻¹), typically detaches before the “ungated” front C-head (expected detachment rate of ∼30 s⁻¹; Fig. 4d).

To examine whether gating occurs for the front C-head in the two-head-bound state, we introduced an E236A point mutation into the N-head (termed N-head/E236A; Fig. 5c), which kinetically traps the head in a microtubule-bound, neck-linker docked state^3,30^. In smFRET observations, the molecules predominantly showed low FRET (Fig. 5d, e), indicating that the C-head mainly occupies the leading position. The ATP turnover rate of this mutant was 25.4 s⁻¹ per chain (Fig. 5f)—about half that of the dimeric kinesin (40 s⁻¹ per head) and is comparable to that of the detachment rate of the ungated rear C-head (Fig. 4d). These findings demonstrate that the front C-head lacks gating regulation and can repeatedly hydrolyze ATP, detach, and rebind to the front binding site. In contrast, when we introduced the E236A mutation into the C-head (termed C-head//E236A), the tandem kinesin predominantly showed high FRET (Fig. 5d, e), indicating that the N-head occupies the leading position, while its ATPase activity was severely suppressed (2.5 s^-1^; Fig. 5f). These results support the front-head gating mechanism in which backward strain on the neck linker suppresses ATP hydrolysis and subsequent detachment of the front head.

## Discussion

The unidirectional and processive movement of tandem kinesin demonstrates that both biased-diffusion and biased-binding mechanisms can drive forward stepping. However, in the two-head-bound state, with the N-head leading and C-head trailing, the detached C- head often rebinds to the rear-binding site. Meanwhile, the leading N-head remains securely bound to the microtubule, gated by its backward-pulled neck linker, until the C-head successfully binds to the forward-binding site. These findings suggest that front-head gating is fundamental for both kinesin-1’s directionality and processivity. As long as front-head gating persists (the maximum detachment rate of the front head for dimer is ∼6 s^-1^)^30^, the tethered head can repeatedly rebind to the rear-binding site without compromising directional and processive movement. In this context, the biased-binding mechanism is not essential for directionality but rather enhances stepping efficiency, that is velocity (under no load) and coupling ratio, by preventing the rebinding events. In contrast, when the C- head leads in the two-head-bound state and front-head gating is not effective, the combination of rear-head gating and biased-binding of the N-head serves as a backup mechanism to maintain processive motility.

The forward stepping rate also depends on how fast the detached head reaches the forward-binding site by Brownian diffusion (“first passage time”)—a process unaffected by the biased-binding mechanism^12^. When a hindering load is applied to dimeric kinesin, either through an optical trap or viscous drag during intracellular transport, the neck linker of the tethered head gets pulled backward. This shifts the head’s center of Brownian diffusion backward, significantly increasing the first passage time (estimated to be about 5 µs and 1 sec under 0 and 6 pN hindering load, respectively^33^). While ATP binds to the microtubule-bound head regardless of its neck linker configuration^30^, hydrolysis does not readily proceed when the neck linker is pulled backward (front-head gating). If the forward stepping rate falls below the head’s detachment rate (∼6 s^-1^ at saturating ATP^30^), the microtubule-bound head will detach before the tethered head steps forward, either terminating the processive run or causing the tethered head to bind, resulting in backward steps or slips^36,40,41^. In this condition, the power stroke, specifically the formation of the cover-neck bundle of the microtubule-bound head, acts as an “elastic” lever, helping to counteract the increased first passage time and thereby enhancing both stall force and velocity under load^24,25^. The neck linker docking might be not well-suited for this role because it exhibits a nearly two-state transition between docked and undocked states, and the free energy change associated with neck linker docking is relatively small^42^. The biased-binding mechanism remains essential for efficiency by preventing the detached head from rebinding, maintaining the high coupling ratio between forward steps and ATP hydrolysis under load^43,44^.

Kinetic asymmetry has become the fundamental mechanism explaining how free energy from catalysis drives unidirectional motion in synthetic molecular motors^6–8^. We recently proposed a kinesin version of kinetic asymmetry termed the tension-based regulation mechanism^26^, in which neck linker strain asymmetrically regulates the rates of allosteric conformational transitions, rather than directly controlling chemical transitions. Unlike synthetic motors where chemical and structural changes occur almost simultaneously, kinesin’s nucleotide-pocket and neck linker base are separated by ∼2.5 nm and communicate through allosteric conformational changes^26,32^. These conformational changes, driven by thermal fluctuations, must overcome significant activation energy barriers that are modulated by neck linker tension^26^. This process typically takes milliseconds and subsequently modulates chemical transition rates (e.g., ATP dissociation and hydrolysis rates differ between open and closed states^30^). The N-head’s preferential binding to forward sites and its ATP-independent behavior (Fig. 3d) can be fully explained by this mechanism. When the tethered N-head binds to the forward- or rearward-binding site before ATP binds to the microtubule-bound C-head, the tandem kinesin transitions into a two-head-bound state (both in nucleotide-free/open state; Fig. 6c); either the N-head leads (termed T_openC_-L_openN_), or the C-head leads (termed T_openN_-L_openC_). The T_openC_-L_openN_ state is thermodynamically more favorable than the T_openN_-L_openC_ state, because in T_openN_-L_openC_, the neck linker of the trailing N-head must navigate around a bulge on the head’s surface to stretch forward (Supplementary Fig. 1b). In contrast, the cover strand of the trailing C- head, which connects to the N-head, projects upward from the base of the neck linker, avoiding interference from this bulge^26^. Consequently, the tethered N-head preferentially binds to the forward site, since this neck linker configuration requires less entropy reduction. This theory also applies to the ATP-dependent microtubule-binding preference of the C-head (Fig. 3d). Prior to ATP binding, the tethered C-head can bind only to the rear-binding site. This is because the neck linker tension is lower in the T_openC_-L_openN_ state (where the C-head binds to the rear site) than in the T_openN_-L_openc_ state (where the C-head binds to the forward site), as explained above. After ATP triggers the conformational change in the N-head, the neck linker tension becomes lower when the C-head binds to the forward site (T_closeN_-L_openC_ state) compared to when it binds to the rear site (T_openC_-L_closeN_ state), enabling the C-head to step forward (Fig. 6d).

**Figure 6.**
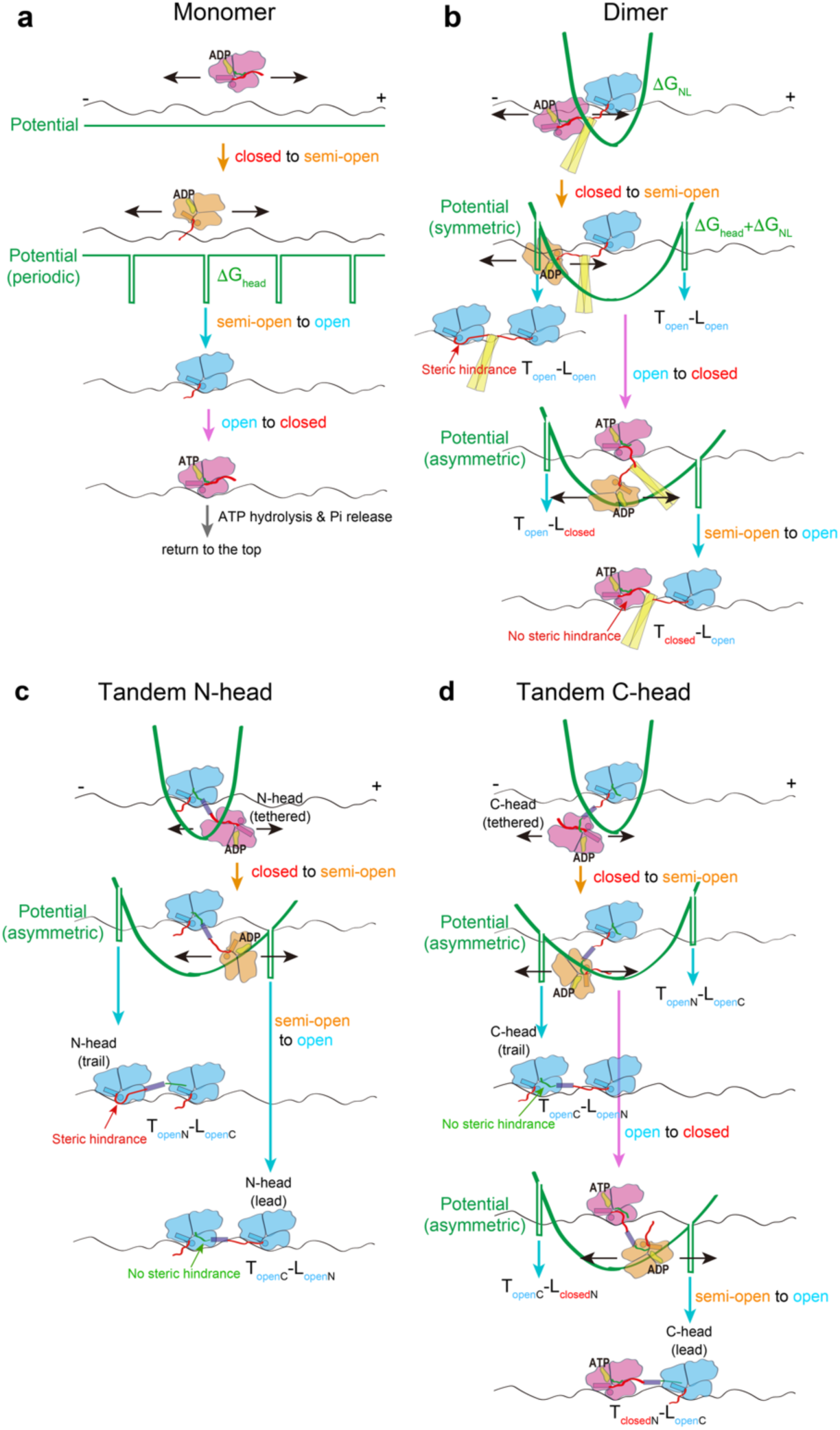
Kinetic asymmetry in conformational transitions determines the binding site preference of the tethered head. Schematic explaining how the microtubule-binding rate of the detached monomer (**a**), tethered head of a dimer (**b**), tethered N-head (**c**) and C-head (**d**) of tandem kinesin are controlled. Three major conformational states (semi-open, open and closed) of kinesin head are indicated in orange, cyan and magenta, respectively. Cover strand and neck liner are depicted in green and red. Potential landscapes along the microtubule-long axis in each conformational state are shown in bold green lines. These potentials consist of two components: the square well potential of microtubule-binding when the head has no neck linker constraint (ΔG_head_), and the harmonic potential that arises from entropy reduction as disordered neck linkers (entropic springs) stretch (ΔG_NL_). ΔG_head_ has a large negative value and remains constant regardless of binding site. In contrast, ΔG_NL_ is positive and becomes asymmetric based on where steric hindrance interferes with neck linker stretching. The tethered head’s binding site preference is determined by differences in neck linker tension between the two two-head-bound state, which creates an asymmetry in the activation energies of microtubule binding. The power stroke modifies the potential energy landscape in a defined area between binding sites and influences the first passage time of the tethered head, yet it does not substantially affect binding site preference. We hypothesized that the detached head initially exists in an ADP-bound/closed state, based on the previous findings that ADP-bound kinesin transition between neck linker docked and undocked states in the absence of microtubules^36^, and that the neck linker docked, ADP-bound state is incapable of microtubule-binding^37^. This mechanism prevents the detached head from rapidly rebinding to its original binding site after dissociation.

Notably, neither ATP hydrolysis nor ATP binding is required for N-head to step forward. The free energy for this step comes from the thermodynamically favorable conformational transitions of microtubule binding and semi-open (ADP-bound, weak microtubule-bound state) to open state (Fig. 6a). The difference in neck linker tension—and its associated entropy reduction—between the two two-head-bound states produces a kinetic asymmetry in the semi-open to open allosteric conformational transition, which suffices for the forward step through thermal fluctuation. In dimeric kinesin, ATP binding to the microtubule-bound head is required because the trailing head must transition to the closed state to reduce neck linker tension in the two-head-bound state (Fig. 6b)^26^. However, the energy from ATP hydrolysis is not necessary for the forward step—rather it primarily functions to detach the tightly bound head from the microtubule. The front and rear head gating mechanisms can also be explained by kinetic asymmetry based on tension-based regulation of conformational transitions between open and closed states as previously described^26^. Both the leading head’s open-to-closed transition and the trailing head’s closed-to-open transition are suppressed by entropy reduction resulting from increased neck linker tension (Supplementary Fig. 16).

## Conclusion

In summary, regulations of both microtubule-binding and detachment of the heads during processive movement of dimeric kinesin-1 can be explained by kinetic asymmetry in allosteric conformational changes. The kinetic asymmetry in microtubule-binding affects unloaded velocity and coupling ratio, while the power stroke, which influences first passage time, enhance stall force and velocity under load. Free energy comes from thermodynamically favorable conformational changes upon microtubule binding, with neck linker tension asymmetrically increasing this energy through entropy reduction. The kinetic asymmetry in microtubule-detachment (especially front-head gating) is essential for kinesin’s directional and processive movement, which is also created by entropy reduction of the stretched neck linker. The neck linker functions as a tension sensor (Supplementary Fig. 16), ensuring that the trailing head preferentially hydrolyzes ATP and detaches from microtubule, a process that consumes most of the ATP hydrolysis energy. This aligns with our energetic study showing that only 20% of the input energy converts to mechanical work, while the remainder dissipates internally^45^ (presumably as information flow^27,46,47^). Future studies of these mechanisms under load will enhance our understanding of how kinesin converts free energy into mechanical work under thermal fluctuations.

## Acknowledgments

We thank M. Nakajima and Y. Sakai for support with cloning. H. Ueno for advice in building high-speed dark-field microscope. R. Vale for insightful comments on the manuscript. M.T. was supported by MEXT KAKENHI Grant Number 17H03661 and by JSPS KAKENHI Grant Numbers JP20H05542, JP22H04850.

## Author contributions

M.T. and H.I. conceived the experiments; H.I. performed single molecule fluorescence and FRET observations and performed ATPase measurement; K.M. performed the dark-field microscopy observations; H.I. and K.M. analyzed the data, and M.T., H.I., K.M. interpreted the results; H.I. and K.M. prepared figures, and M.T. wrote the manuscript.

## ONLINE METHODS

### DNA cloning and protein purification

Tandem kinesin was constructed by joining two 336 amino-acid “cysteine-light” human kinesin-1 monomer (K336CLM) sequence in tandem. The two K336CLM were connected using a poly-Pro linker of varying length (Supplementary Fig. 1a). A Glycine residue flanked the poly-Pro linker and the first K336CLM (N-head), while GT sequence (KpnI cloning site) flanked the second K336CLM (C-head) and the poly-Pro linker. A GT sequence (second KpnI site for cloning) and His_6_-tag were added to the C-terminus of the C-head. E215C and S43C were introduced for single-molecule fluorescent and smFRET observations, and S55C into N- or C-head was introduced for high-speed dark-field microscopy (position of introduced cysteine for linker-labeling is shown in Supplementary Fig. 8a). 490 amino-acids cysteine-light homodimer or heterodimer kinesin was used as a wild-type dimer^39,48^. Cysteines and mutagenesis (E236A) were introduced by PCR cloning involving QuikChange mutagenesis (Stratagene)^48^. All the constructs were verified by DNA sequencing. The DNA containing the C-head was digested with KpnI and subcloned into the pET17b vector carrying the N-head to generate a tandem kinesin construct. Kinesin proteins were expressed and purified as described previously^39,48^. Heterodimeric kinesins were expressed in E. coli BL21(DE3), purified by two-step affinity chromatography for nicket-nitrilotriacetic acid and Strep-Tactin and dialyzed as described previously^39^. Tandem kinesins were expressed in E. coli BL21(DE3), purified by affinity chromatography for nicket-nitrilotriacetic acid and dialyzed as described previously^48^. Tubulin was purified and polymerized as described previously^49^. Protein concentrations were determined by Bradford assays using BSA as a standard. For single-molecule fluorescent and FRET observations, dialyzed proteins were reacted with Cy3-maleimide (Invitrogen PA23031) and Cy5- maleimide (Invitrogen PA25031) at a motor/Cy3/Cy5 molar ratio of 1:10:10, for 10 min at room temperature. Unreacted dyes were quenched with 1 mM DTT and removed through microtubule affinity purification as described^48^.

### ATPase measurement

Microtubule-stimulated ATPase activity was measured using spectrometer (V-550; JASCO) with a couple enzymatic assay as describe previously^49^. The ATPase assays were performed in BRB12 buffer (12 mM PIPES (pH 6.8), 1 mM MgCl_2_, 2 mM EGTA) containing 1 mM ATP, 20 μM taxol, 2 mM DTT, 50 mM NaCl, 0.1 mg/ml casein and coupled NADH oxidation system (0.2 mM NADH, 5 mM phospho(enol)pyruvate, 10 μg/ml of pyrvate kinase, and 10 μg/ml of lactate dehydrogenase). The assays were started with 10 nM kinesin and varying concentrations of microtubules at 22°C.

### Single molecule fluorescence observation

Individual Cy3/Cy5-labeled tandem kinesin moving along axonemes (purified from sea urchin sperm flagella) were observed in a custom-build prism-type laser-illuminated total-internal-reflection microscope as described^48^. Assay chamber constructed between a quartz slide and a coverslip was first filled with axonemes diluted in BRB12 for 3min, followed by a washing with 1mg/ml casein in BRB12 for 3 min. The chamber was then filled with the assay solution contained Cy3/Cy5-labeled kinesin, 1 mM ATP, 70 mM 2- mercaptoethanol and oxygen scavenging system (4.5 mg/ml glucose, 50 U/ml catalase, 50 U/ml glucose-oxidase) and sealed. Cy5 fluorophores were exited at 635 nm (Radius 635, Coherent) and fluorescence was recorded at 10 ms temporal resolution. Images were analyzed using Image J, and velocities and run lengths were determined from kymographs.

### Single-molecule FRET observations

Single molecule FRET measurements were carried out using a custom-built prism-type total internal reflection fluorescence microscopy as described^18,39^. The chamber was constructed as described above, except that an ATP regenerating system was added to the final solution^39^. Dual-labeled molecules bound to the axoneme in the presence of 5-1000 µM ATP, 1 mM AMP-PNP or 200 nM ADP were excited with 514 nm argon laser (35LAP321; Melles Griot), and donor and acceptor fluorophores were simultaneously observed using a Dual-View (Optical Insights) and EM-CCD camera (iXon DV860; Andor) as previously described^18^ at acquisition rates of 50 frames/s for the static FRET observations using AMP-PNP and ADP) and 100 frames/s for the dynamic FRET observations in the presence of ATP. Images were analyzed with Image J with custom-designed plug-in software as described previously^18^.

### Gold labeling of kinesin

Purified and dialyzed kinesin was incubated with biotin-PEG2-maleimide (Thermo Fisher 21901) at a molar ratio of 1:80 (peptide:biotin) for 10 min at room temperature and unreacted biotin-PEG2-maleimide was deactivated by adding DTT (10 mM final concentration). Biotinylated kinesin was further purified by microtubule affinity purification as described previously^21^. Biotinylated kinesin and microtubule were mixed in the presence of 1 mM AMP-PNP and 50 µM taxol and incubated for 15 min at room temperature. Microtubule-bound kinesin was then collected by centrifugation (230,000*g* for 10 min) and was added streptavidin solution containing 12 mM PIPES (pH6.8), 2 mM MgCl2, 1 mM EGTA, 20 μM taxol and 2.5 mg/ml streptavidin (191-12851; Wako Pure Chemical Industries) and incubated 20 min at room temperature. Microtubule-bound streptavidin-modified kinesin was collected by centrifugation (230,000g for 10 min) and released from microtubules by adding BRB12 buffer containing 100 µM ATP and 200 mM KCl as described previously^21^. Biotin-coated gold nanoparticles were prepared as previously described^21^ except streptavidin was not added. In brief, colloidal gold particles of 40 nm in diameter (EM.GC40, Boston Biomedical) was reacted with an alkanethiol solution containing 0.01% biotin-EG6-undechanethiol (SPT-0012D, SensoPath Technologies), 0.1% carboxy-EG6-undecanethiol (C445-12, Dojindo Laboratories) and 0.1% hydroxy-EG6-undecanethiol (C355-12; Dojindo Laboratories) and incubated overnight at 70°C. Unreacted alkanethiol was removed by centrifugation (20,000*g* for 10 min by 5 times) and resuspended in 10 mM Boric-acid buffer (pH 8.0). Just before observation, streptavidin-modified kinesin and biotin-coated gold nanoparticles were mixed at a 1:1 molecule/particle ratio for 15 min on ice.

### Total internal dark-field microscopy

A flow cell was constructed and microtubules were attached on the glass surface by using protein A and anti-α-tubulin antibody as described previously^21,30^. The cell was then infused with 40 μl of BRB 12 buffer (12 mM PIPES (pH 6.8), 2 mM EGTA and 1 mM MgCl2) containing 20 μM taxol and 1 mg/ml casein for 2 min. The cell was infused with BRB 12 buffer containing 3 nM gold-labeled kinesin for 5 min, and unbound kinesins and gold nanoparticles were removed by washing with 60 μl of the BRB 12 buffer containing 20 μM taxol. Finally, the flow cell was infused with 40 μl of the observation buffer (BRB 12 buffer containing 70 mM β-mercaptoethanol, 10 μg/ml creatine kinase, 2 mM creatine phosphate, 20 mM KCl and 5-1000 µM ATP), sealed with nail polish and used for observation. All procedure were carried out at room temperature. The gold-labeled kinesin was observed using the total internal reflection dark-field microscope as described previously^21^ and the scattering images were recorded with a high-speed CMOS camera (FASTCAM Mini AX100; Photron) at a frame rate of 20,000 fps (50 µs temporal resolution). Observations were carried out at 24-26°C.

### Data analysis for the dark-field observations

The coordinates of gold nanoparticles in raw dark-field images (Xraw, Yraw) were determined frame by frame by fitting the intensity profiles with a two-dimensional Gaussian function using the Levenberg-Marquardt algorithm with JAVA library Apache Commons Math 2.2 software^21,30^. The ‘bound’ and ‘unbound’ states of gold-labeled kinesin heads were determined as follows: two-dimensional standard deviation of the trajectory for each time frame t was calculated as [t - 20, t + 20] and the median μ and standard deviation (s.d.) σ of the s.d. of the trajectory was determined by fitting the histogram of the s.d. with a Gaussian function. Transitions from the bound to unbound state were detected when the s.d. was higher than μ+ 4σ in 10 consecutive frames and transition from the unbound to bound state were detected when the s.d. was lower than μ+ 4σ in 10 consecutive frames. The accurate frame of the transition was determined by fitting the on- and off-axis trajectory near the transition with two state step function. The rate constant of each state was determined by fitting the histogram of the dwell time with single exponential functions.

