## Supplemental Figures for "Kinetic asymmetry drives kinesin-1’s unidirectional and processive movement"

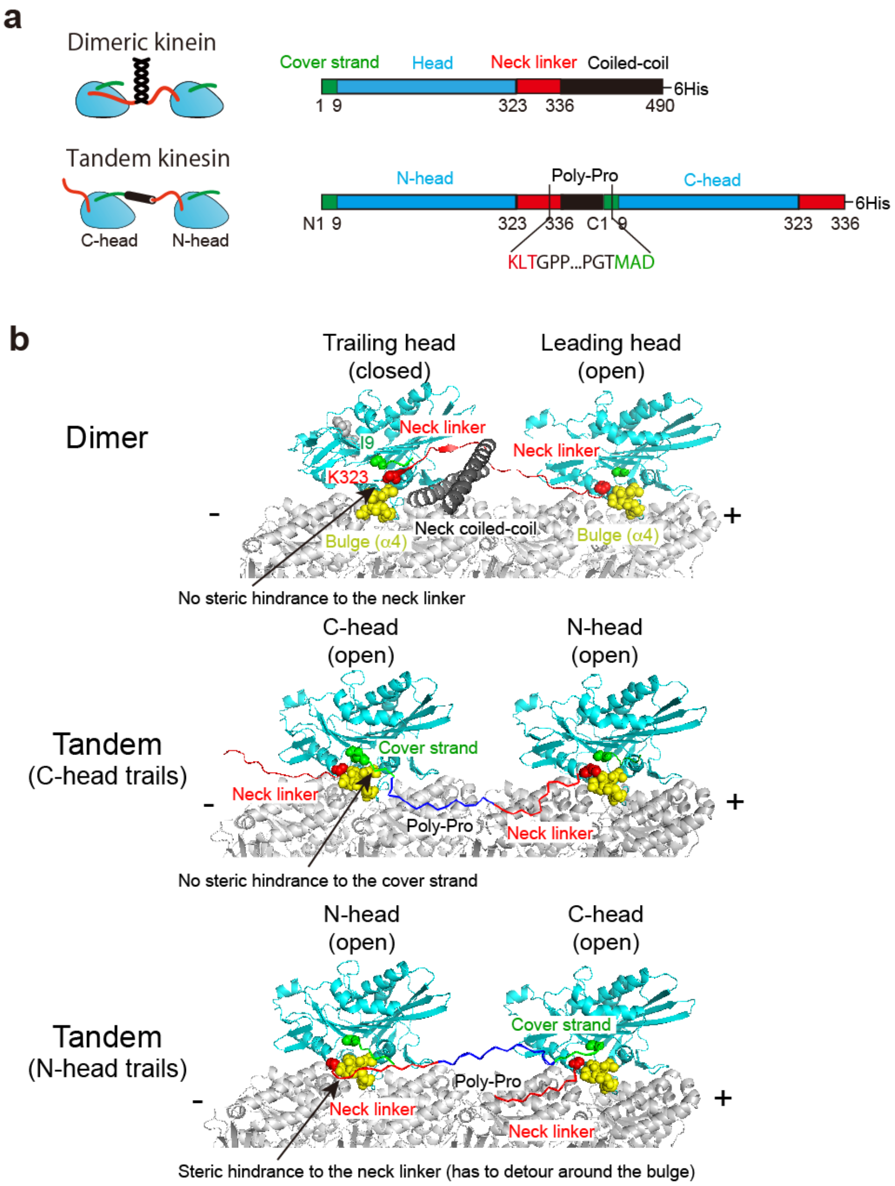


**Supplementary Figure 1 | Design and structure of tandemly-joined two-headed kinesin (tandem kinesin) compared with dimeric kinesin.** (**a**) Polypeptide designs of dimeric kinesin (490 amino acids) and tandem kinesin, which consists of two 336-amino-acid heads connected by poly-proline inserts of varying length. The poly-Pro sequence was flanked by glycine and GT (KpnI cloning site) sequences, which served as flexible joints. The N-terminal cover strand and C-terminal neck linker of the head are shown in green and red, respectively. Key residue numbers are indicated. (**b**) Atomic models of dimeric kinesin and tandem kinesin on the microtubule in two-head-bound states (side-view). Both heads of tandem kinesin are nucleotide-free/open state (prior to ATP binding). The base residues where the cover strand and neck linker emerge from the head are shown in space-filling (I9 and K323, respectively). The C-terminus of α4 helix (L268-S272), which acts as a bulge that interferes with the neck linker in the open state when stretched forward[1], is shown in space-filling (yellow). This bulge does not interfere with the cover strand of the trailing C-head in the open state when stretched forward, as the base of the cover strand (I9) is positioned above the bulge.

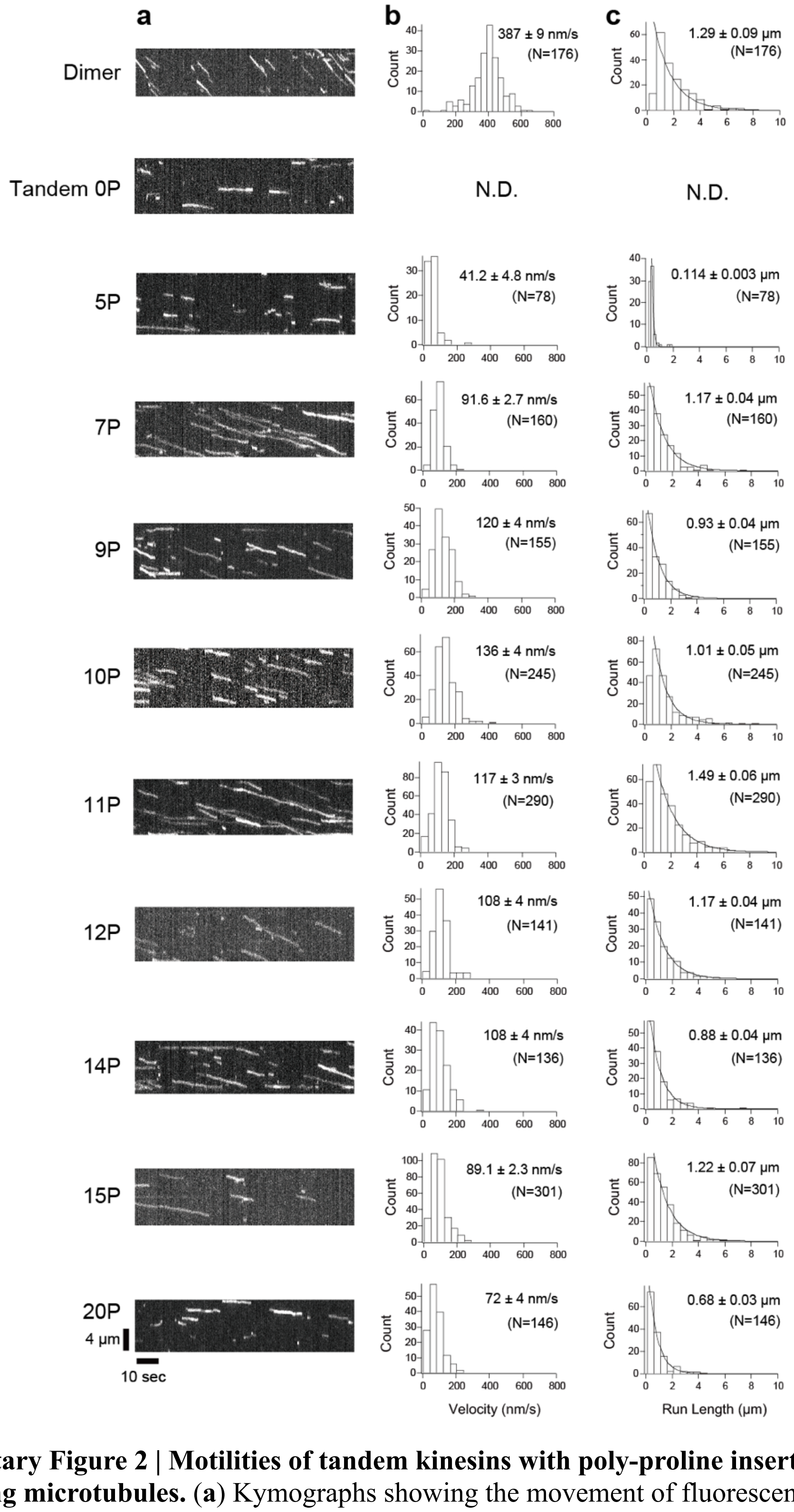


**Supplementary Figure 2 | Motilities of tandem kinesins with poly-proline inserts of various lengths along microtubules.** (**a**) Kymographs showing the movement of fluorescently labeled tandem kinesins with varying linker size and dimeric kinesin along axonemes. (**b, c**) Histograms showing the distributions of velocity (**b**) and run length (**c**). Lines show exponential fit. Numbers indicate mean velocities and run lengths (± s.e.m.). Numbers in parentheses indicate the number of molecules analyzed.

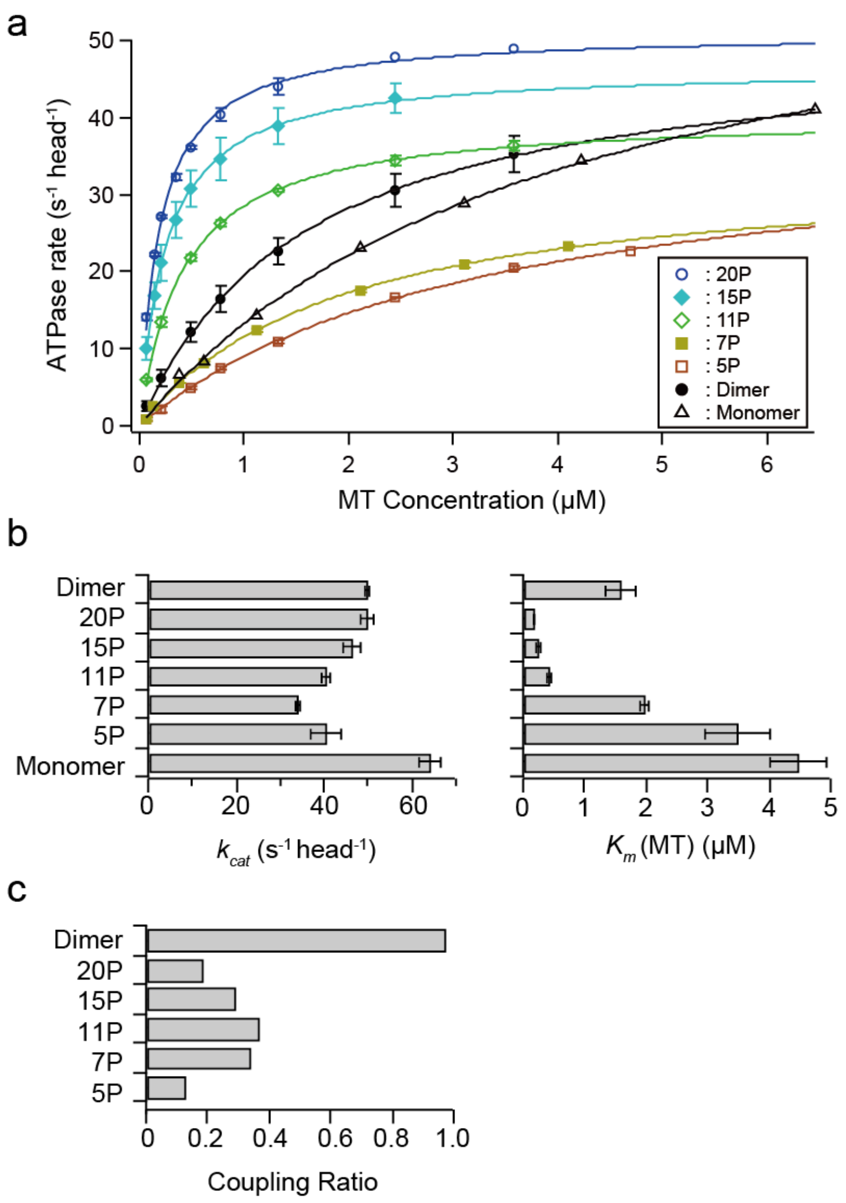


**Supplementary Figure 3 | Microtubule-activated ATP turnover rates of tandem kinesins.** (**a**) Microtubule-activated ATPase rates (per head) for tandem kinesins with various poly-Pro linker sizes, along with dimeric (490 amino-acids) and monomeric (349 amino-acids) kinesins. The rates were plotted against microtubule (MT) concentrations (N = 3; average ± s.e.m.), with solid lines showing the Michaelis-Menten equation fit. (**b**) $k_{cat}$ and $K_m$(MT) values obtained from the Michaelis-Menten fit shown in **a**. (**c**) The coupling ratio calculated by dividing mean velocity (Supplementary Fig. 2) by $k_{cat}$ and 8 nm. This value represents the probability of an 8 nm step per ATP hydrolysis for each dimer or tandem molecule.

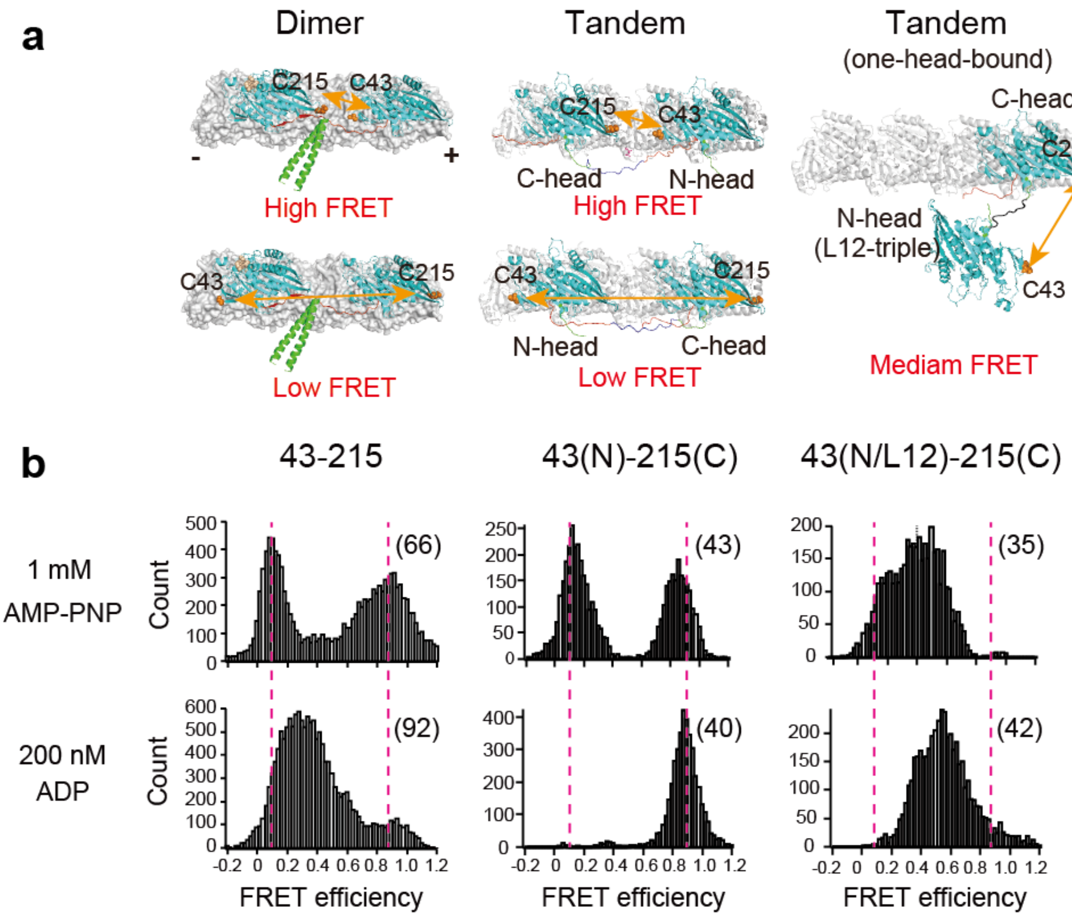


**Supplementary Figure 4 | SmFRET observations of dimeric and tandem kinesins in two- and one-head-bound states on microtubules.** (**a**) Diagrams showing the positions of cysteine residues for dye labeling (C215 and C43; orange spheres) in the two- or one-head-bound states of dimeric and tandem kinesins. (**b**) Histograms of FRET efficiencies of axoneme-bound wild-type dimer, tandem kinesin, and tandem kinesin with L12-triple mutation (that cannot bind microtubules[2,3]) into N-head. Parentheses show the numbers of molecules analyzed. Red dotted lines highlight peaks characteristic of putative two-head-bound state (90 and 10% FRET efficiencies[2]). Under low ADP condition (mimicking ATP waiting state), tandem kinesin showed a single peak at high FRET, representing the two-head-bound state where the N-head leads and the C-head trails. This finding contrasts with the dimer which showed one-head-bound state under low ADP condition, and suggests that tandem kinesin can adopt a two-head-bound state in which N-head leads and both heads are nucleotide-free/open state (Supplementary Fig. 1b).

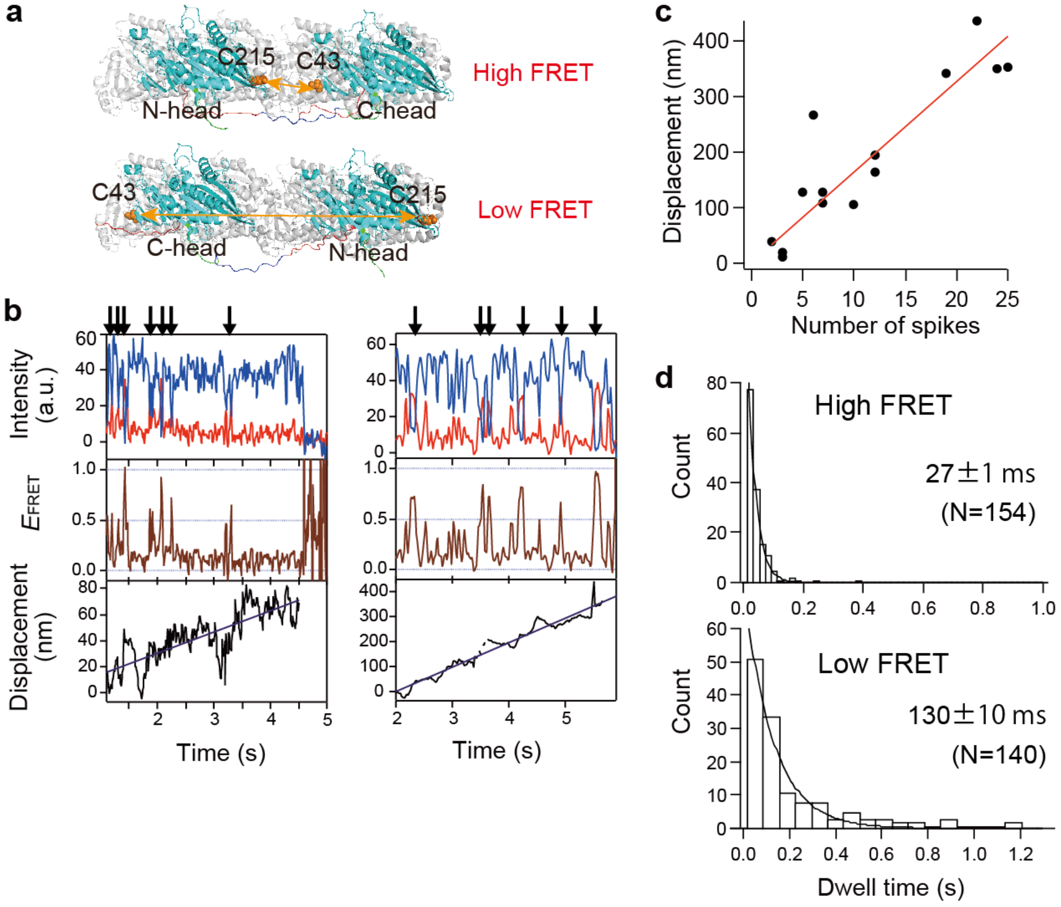


**Supplementary Figure 5 | SmFRET observation of head-head configuration of tandem kinesin with opposite dye-labeling positions.** (**a**) Diagrams illustrating the positions of cysteine residues for dye labeling: C215 on the N-head and C43 on the C-head (reversed from the labeling positions shown in Fig. 2 and Supplementary Fig. 4). (**b**) Representative traces of fluorescence intensities of donor (Cy3, blue) and acceptor (Cy5, red) fluorophores, FRET efficiency and axial displacement for Cy3/Cy5-labeled tandem kinesin at 1 mM ATP. Arrows indicate transitions to high FRET states. (**c**) Relationship between number of transitions towards the high FRET state and the displacement per observation time; each point represents a different molecule. The solid line shows a linear fit (16.3 nm per high FRET transition). (**d**) Distributions of dwell times in the high and low FRET states. Lines show exponential fit. These results further confirm the finding in Fig. 2 that the C-head steps more slowly than the N-head.

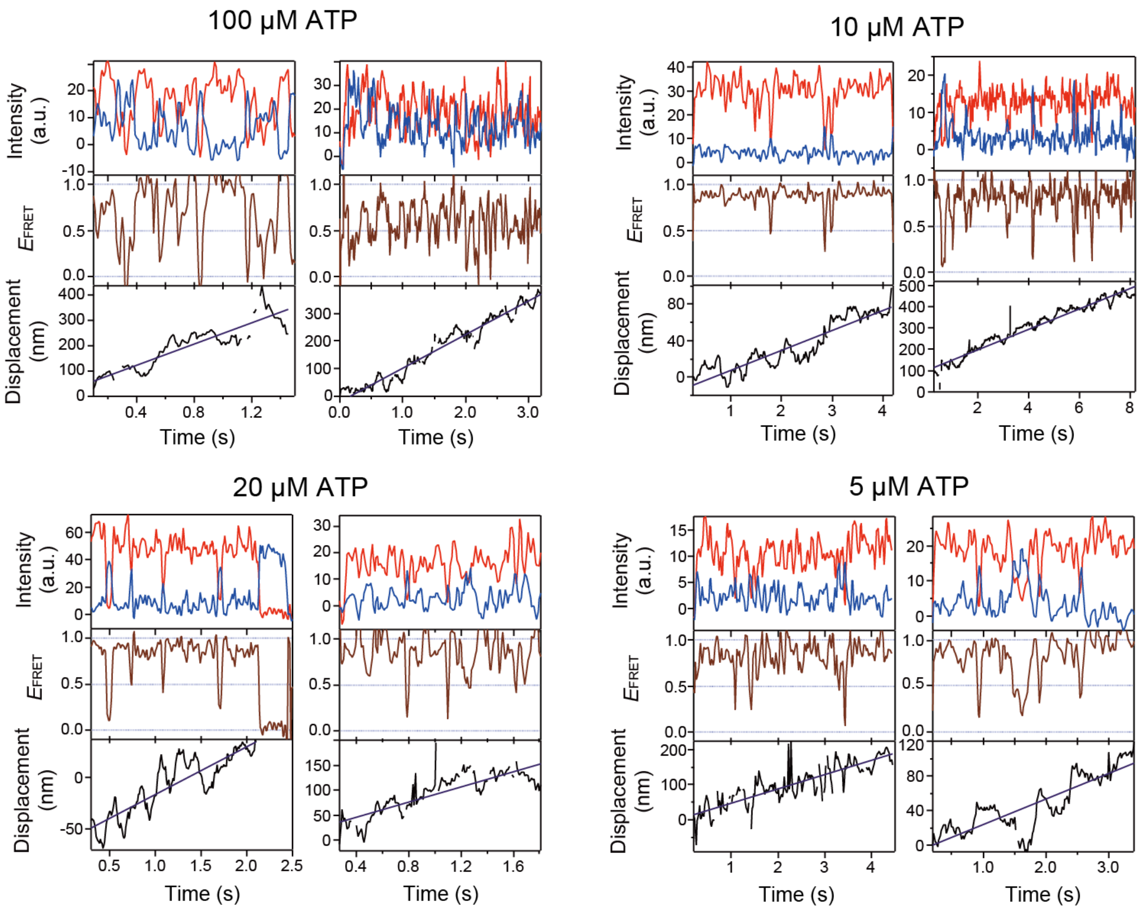


**Supplementary Figure 6 | SmFRET observations of head-head configuration of tandem kinesin under various ATP concentrations.** Representative traces showing fluorescence intensities of donor (Cy3, blue) and acceptor (Cy5, red) fluorophores, FRET efficiency, and axial displacement (with linear fits) for 11P tandem kinesin (C43 in N-head and C215 in C-head; 1 mM ATP traces are shown in Fig. 2b) moving along axonemes at ATP concentrations ranging from 100 to 5 μM.

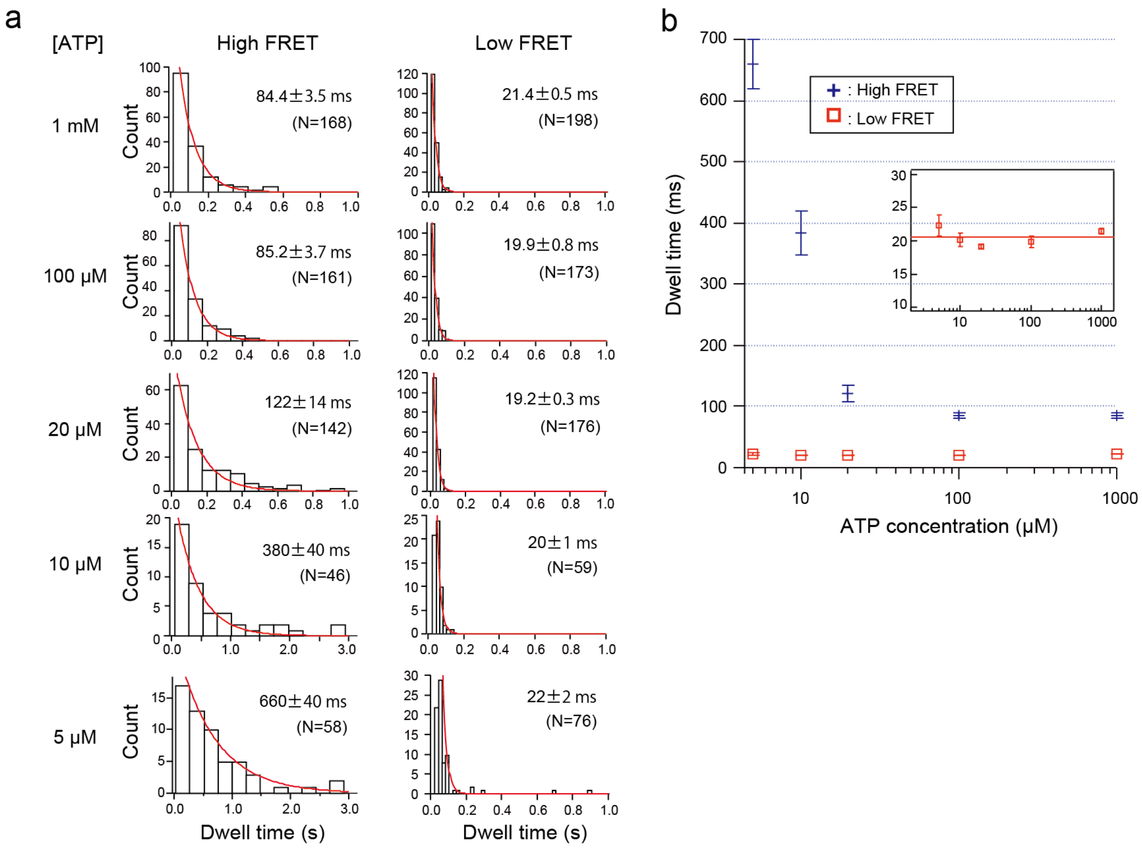


**Supplementary Figure 7 | ATP concentration dependence of dwell times in high and low FRET states of tandem kinesin.** (**a**) Histograms showing the distributions of dwell times in high and low FRET states under various ATP conditions (typical traces are shown in Fig. 2b and Supplementary Fig. 6). Red lines show exponential fit. Numbers indicate mean dwell times (± s.e.m.), with number of dwells analyzed shown in parentheses. (**b**) Mean dwell times of high and low FRET states plotted against ATP concentrations. Red line indicates the average dwell time in low FRET state (20.6 ms), which likely overestimates the actual duration due to limited temporal resolution of smFRET observation (10 ms) (c.f. Supplementary Fig. 13c).

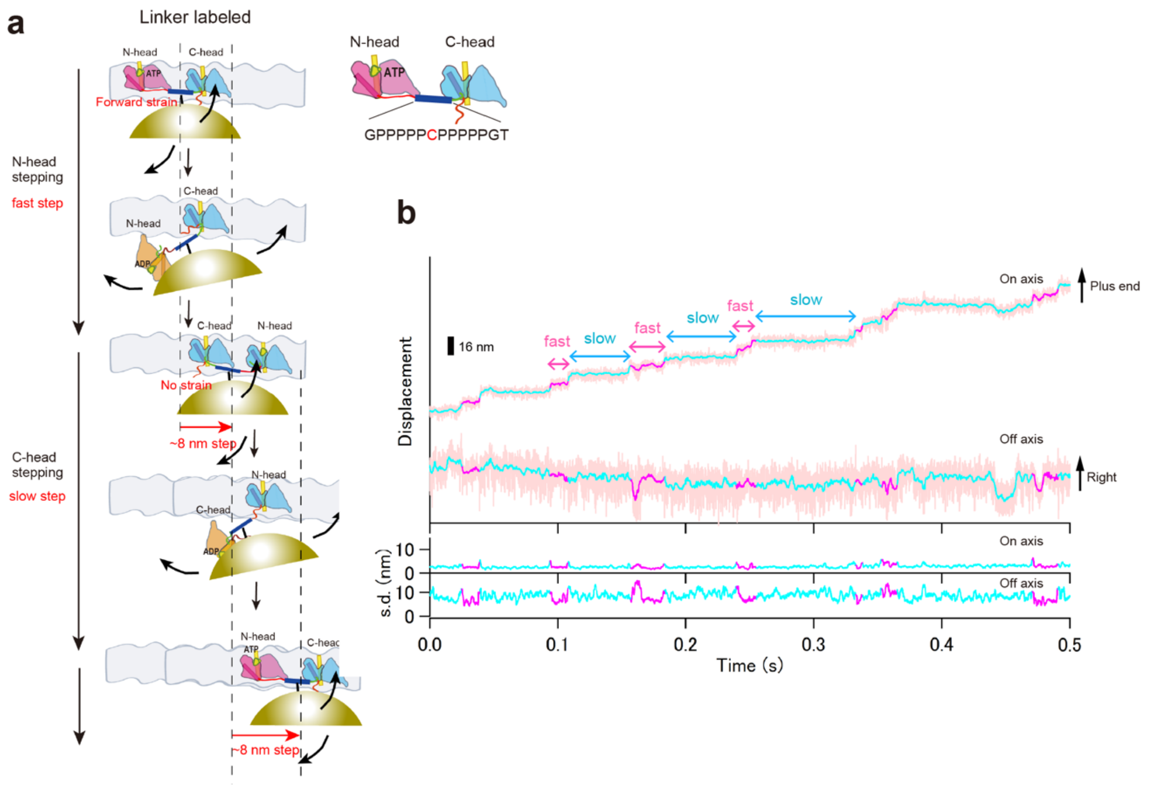


**Supplementary Figure 8 | High-speed single-molecule observation of the gold probe attached to the poly-Pro linker of tandem kinesin.** (**a**) Schematic showing the gold probe attached to the middle of the poly-Pro linker of tandem kinesin moving along microtubules. The middle proline residue of 11P tandem kinesin was replaced with cysteine and was labeled with gold probe. (**b**) Typical trace for the on- and off-axis positions, and standard deviation (s.d.) of the gold probe attached to the poly-Pro linker of tandem kinesin moving along a microtubule in the presence of 1 mM ATP. The gold probe exhibited alternating fast and slow 8-nm steps (magenta and cyan, respectively), representing the alternating N- and C-head stepping. These results are consistent with the smFRET observations that showed alternating high and low FRET states (Fig. 2), indicating that the fast and slow steps correspond to N- and C-head stepping, respectively. The on-axis s.d. was significantly smaller than that in the off-axis direction. This is likely because the linker is stretched in the on-axis direction but has rotational freedom in the off-axis direction during two-head-bound state.

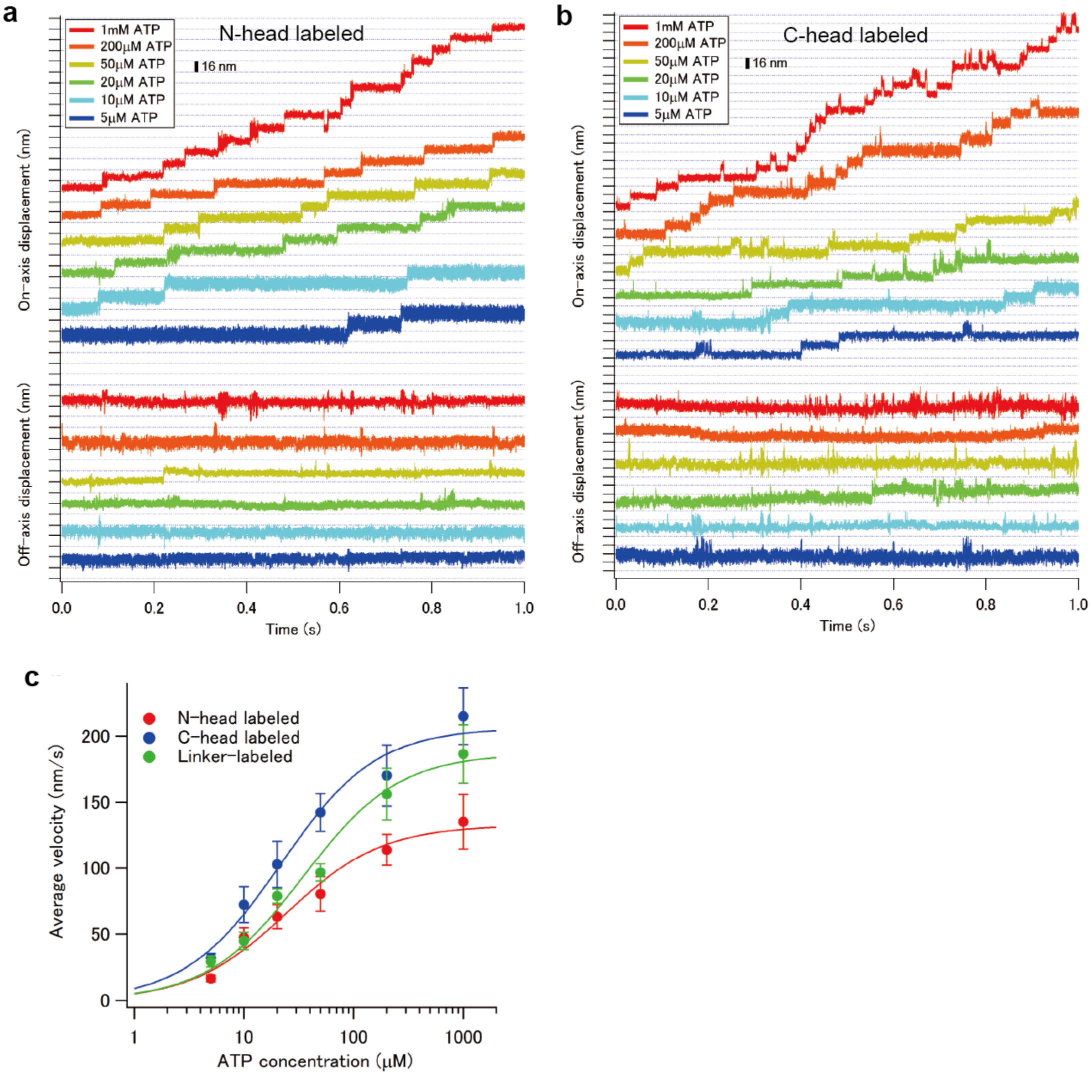


**Supplementary Figure 9 | High-speed single-molecule observations of N- and C-head-labeled tandem kinesin at various ATP concentrations.** (**a, b**) Typical traces of the on- and off-axis positions of the gold probe attached to the N-head (**a**) or C-head (**b**) of 11P tandem kinesin moving along microtubules. (**c**) Average velocities of N-head, C-head and linker-labeled tandem kinesin plotted as a function of ATP concentration. The solid lines show the fit with the Michaelis-Menten equation. The fit parameters were $V_{max}$ = 135 ± 12 nm/s and $K_m$(ATP) 30 ± 5 μM for N-head-labeled; $V_{max}$ = 214 ± 17 nm/s and $K_m$(ATP) 27 ± 4 μM for C-head-labeled; and $V_{max}$ = 187 ± 10 nm/s and $K_m$(ATP) 35 ± 6 μM for linker-labeled tandem kinesins. The maximum velocity of the C-head-labeled tandem kinesin was comparable to that of the linker-labeled tandem kinesin, but that of N-head-labeled tandem was smaller. Since the unbound dwell time of the labeled N-head (~3 ms) was comparable to that of the C-head (Fig. 4b), the reduced velocity of N-head-label results from the extended dwell time of microtubule-bound state of the labeled N-head (Fig. 4c, $\tau_{bound}$). As the measured $\tau_{bound}$ of labeled N-head is an overestimate, we did not use this value for further analysis; instead, we used the short dwell time of the linker-labeled tandem to estimate the detachment rate of N-head. The extended $\tau_{bound}$ of labeled N-head likely occurs because the gold probe on the microtubule-bound N-head interferes with the unlabeled C-head's diffusional search. This interference increases the C-head's rebinding frequency compared to the gold-labeled C-head, which experiences less steric hindrance from the microtubule-bound N-head (without gold labeling).

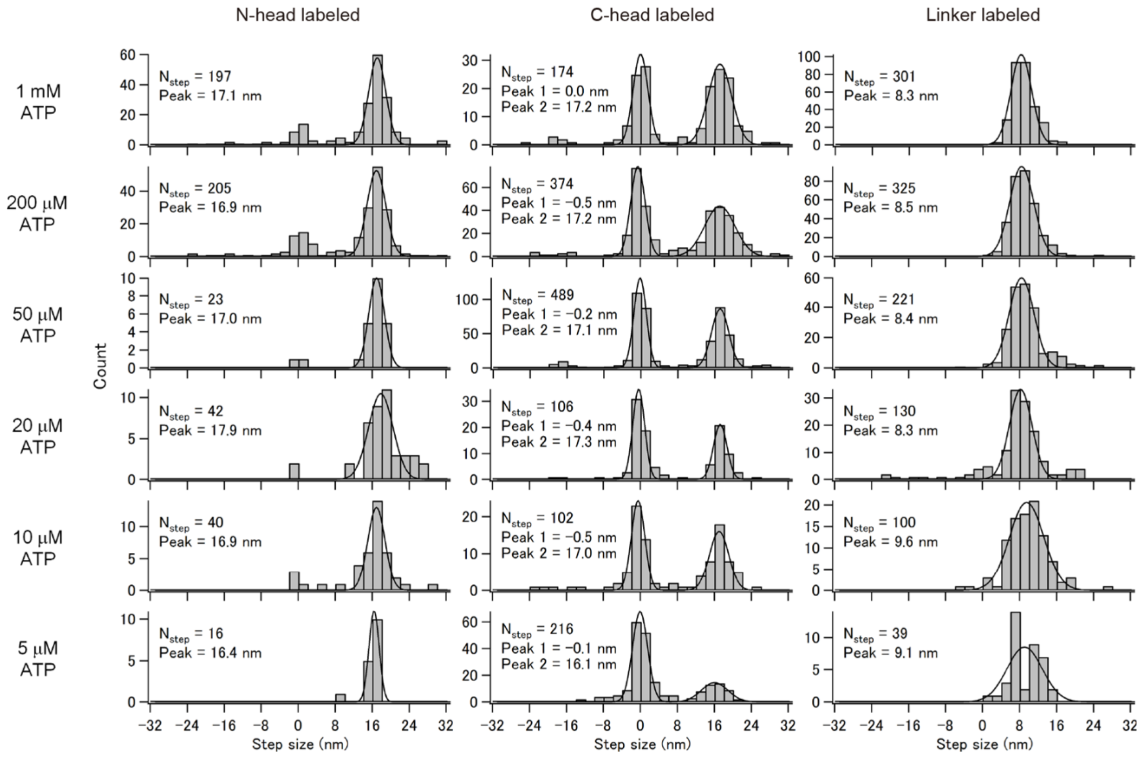


**Supplementary Figure 10 | Step size distributions of the gold-labeled tandem kinesins at various ATP conditions.** Histograms of the step size of gold probe attached to a N-head (left), C-head (middle), and poly-Pro linker (right) at various ATP conditions. The average step sizes determined by fitting with Gaussian distributions (solid lines) are shown.

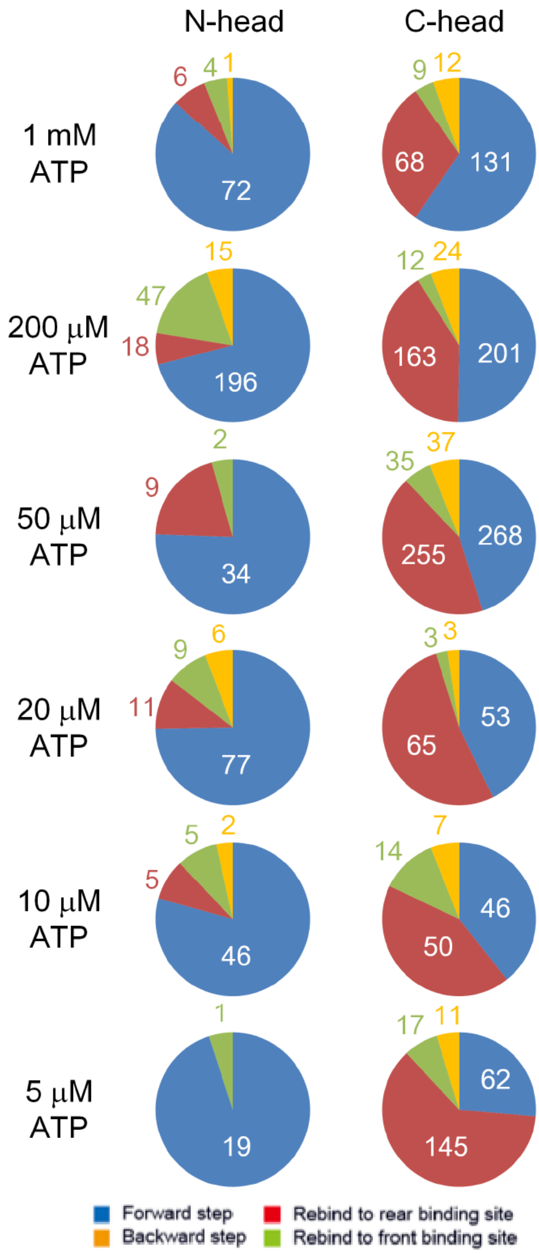


**Supplementary Figure 11 | Classification of the microtubule-detachment and binding events of N-or C-head of tandem kinesin.** Pie-chart showing the number of the binding events observed for N- and C-head labeled tandem kinesin during processive motility at various ATP conditions. We classified the detachment and subsequent binding of the head of tandem kinesin into four types[4]; 1) the trailing head detaches and binds to the forward tubulin-binding site (which leads to the 16-nm forward step of the head, blue), 2) the trailing head detaches and rebinds to the rear tubulin-binding site (the preceding binding site where the head bound before detachment (resulting in 0-nm step, red), 3) the leading head detaches (accompanied with slight displacement of the unbound head toward minus-end of the microtubule) and rebinds to the forward binding site (the preceding binding site (i.e., 0-nm step), green), and 4) the leading head detaches and binds to the rear tubulin-binding site (leading to 16-nm backward step of the head, yellow).

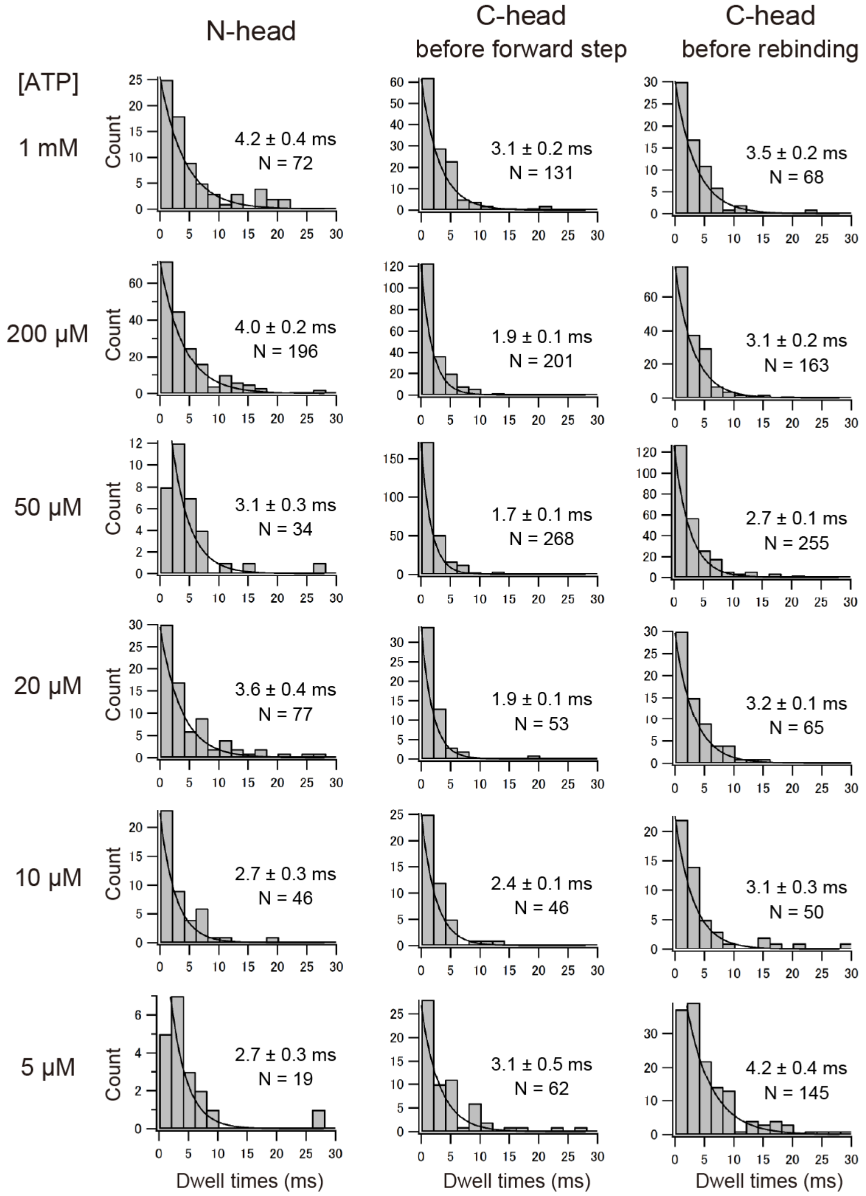


**Supplementary Figure 12 | Distributions of the dwell time in the unbound state of N- and C-heads of tandem kinesin.** Histograms of the dwell time in the unbound states of gold-labeled N- and C-heads of tandem kinesin moving along microtubules at various ATP concentrations. Lines indicate the fit with exponential decay. The average dwell times (± s.e.m.) are shown. The dwell time data of the C-head were separated into two categories: events before forward step and events before rebinding (see Fig. 4a).

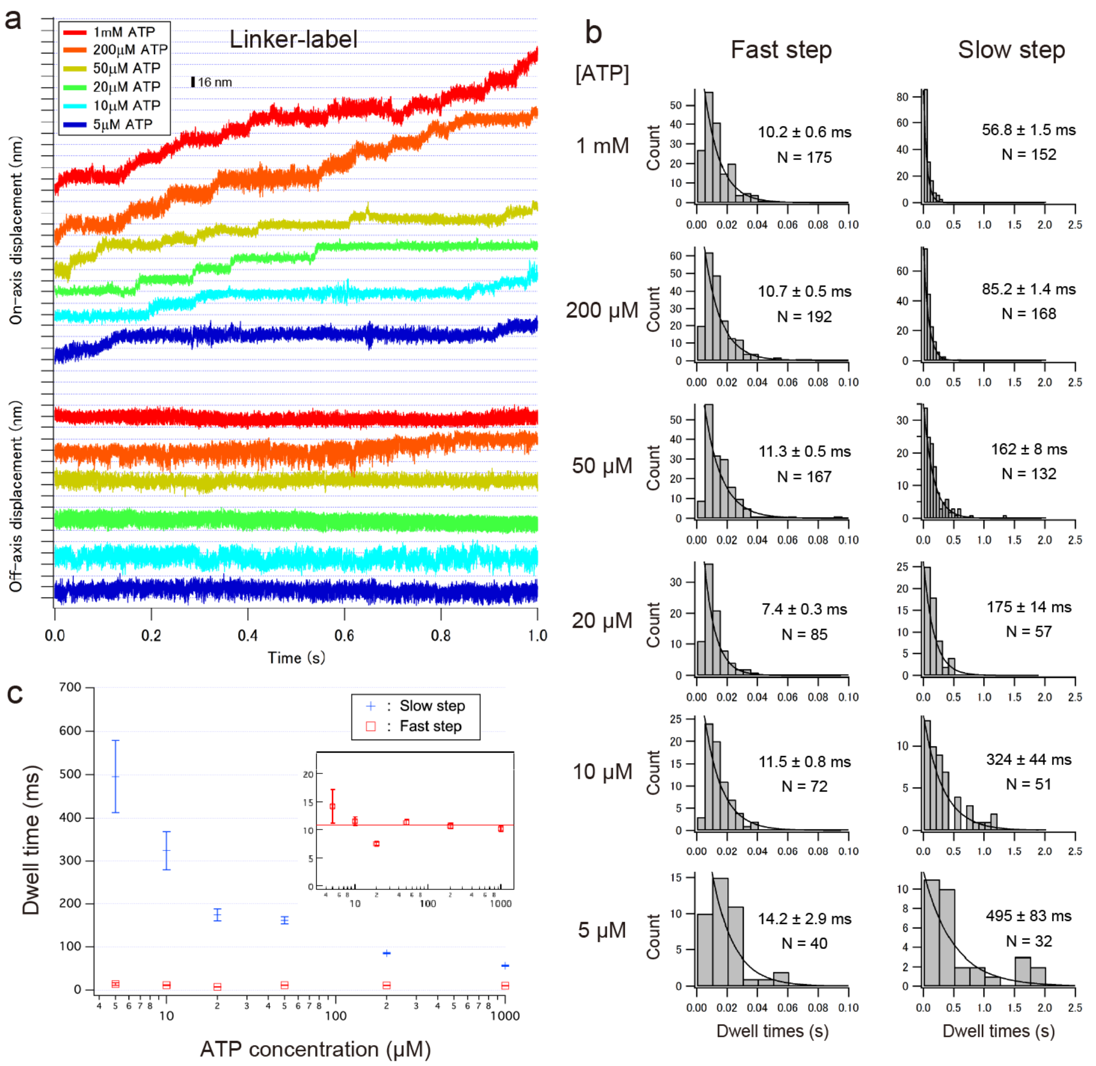


**Supplementary Figure 13 | High-speed dark-field observation of linker-labeled tandem kinesin at various ATP conditions.** (**a**) Typical traces of the on- and off-axis positions of the gold probe attached to the poly-Pro linker of 11P tandem kinesin moving along microtubules at various ATP concentrations. Gold probe attached to the linker of tandem kinesin showed alternating fast and slow steps 8-nm steps (Supplementary Fig. 8). (**b**) Histograms of the dwell time before the fast step and before the slow step. Lines indicate the fit with exponential decay. The average dwell times (± s.e.m.) are shown. (**c**) Mean dwell times before slow and fast steps are plotted against ATP concentrations. The red line indicates the average dwell time before the fast step (10.9 ms), which is more accurate than the measurement obtained using smFRET (20.6 ms; Supplementary Fig. 7b).

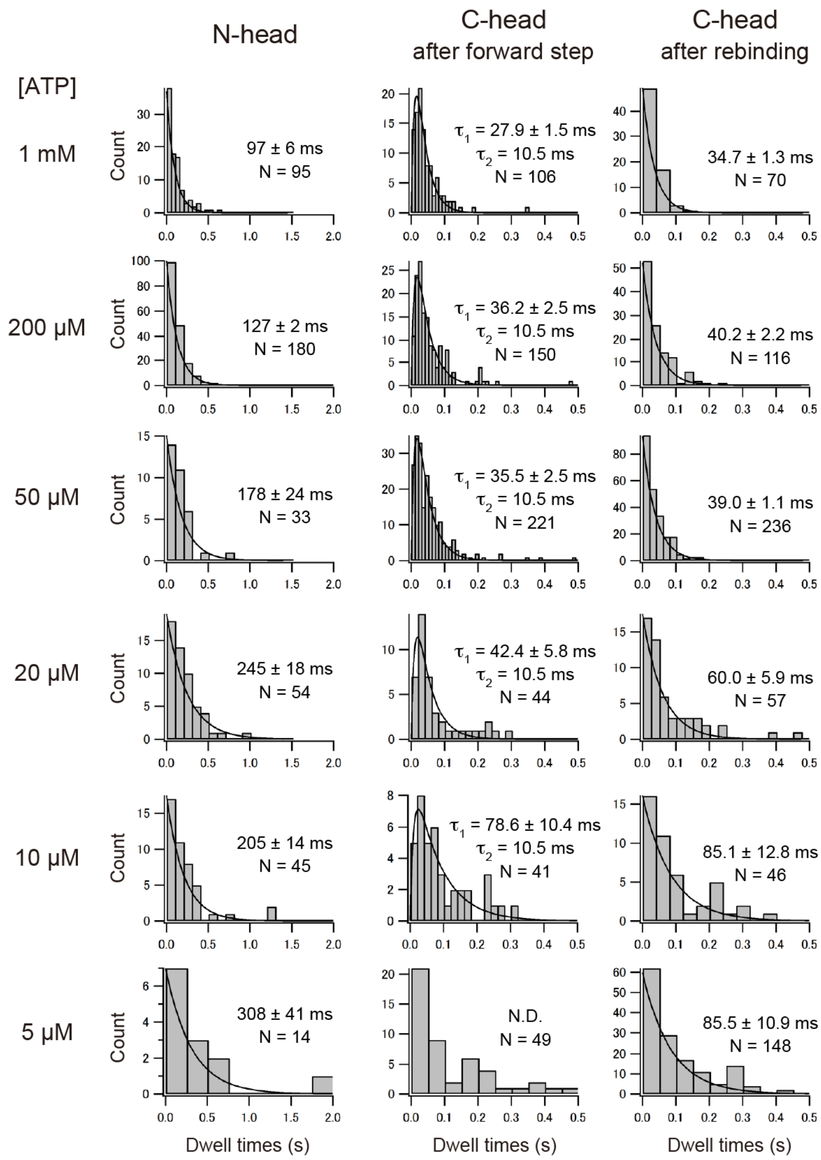


**Supplementary Figure 14 | Distributions of the dwell time in the bound state of the N- and C-heads of tandem kinesin.** Histograms of the dwell time in the bound states of gold-labeled N- and C-heads of tandem kinesin moving along microtubules at various ATP concentrations. Lines indicate the fit with exponential decay, and the average dwell times (± s.e.m.) are shown. The dwell time data of the C-head were separated into two categories: events after forward step and events after rebinding (see Fig. 4a). The histograms of C-head dwell times after forward step fit better with a convolution of two exponentials, with one rate fixed at 10.5 ms (the dwell time of trailing N-head estimated by linker-labeled tandem (Supplementary Fig. 13c)), as the dwell time includes both periods of trailing N-head and trailing C-head (Fig. 4a). The histogram of 5 µM ATP could not be fitted well with a double exponential due to the small sample size. As noted in the legend of Supplementary Fig. 9, the measured dwell time of the labeled N-head is likely an overestimate (due to suppression of biased-diffusional movement of the unlabeled C-head) and was therefore excluded from further analysis.

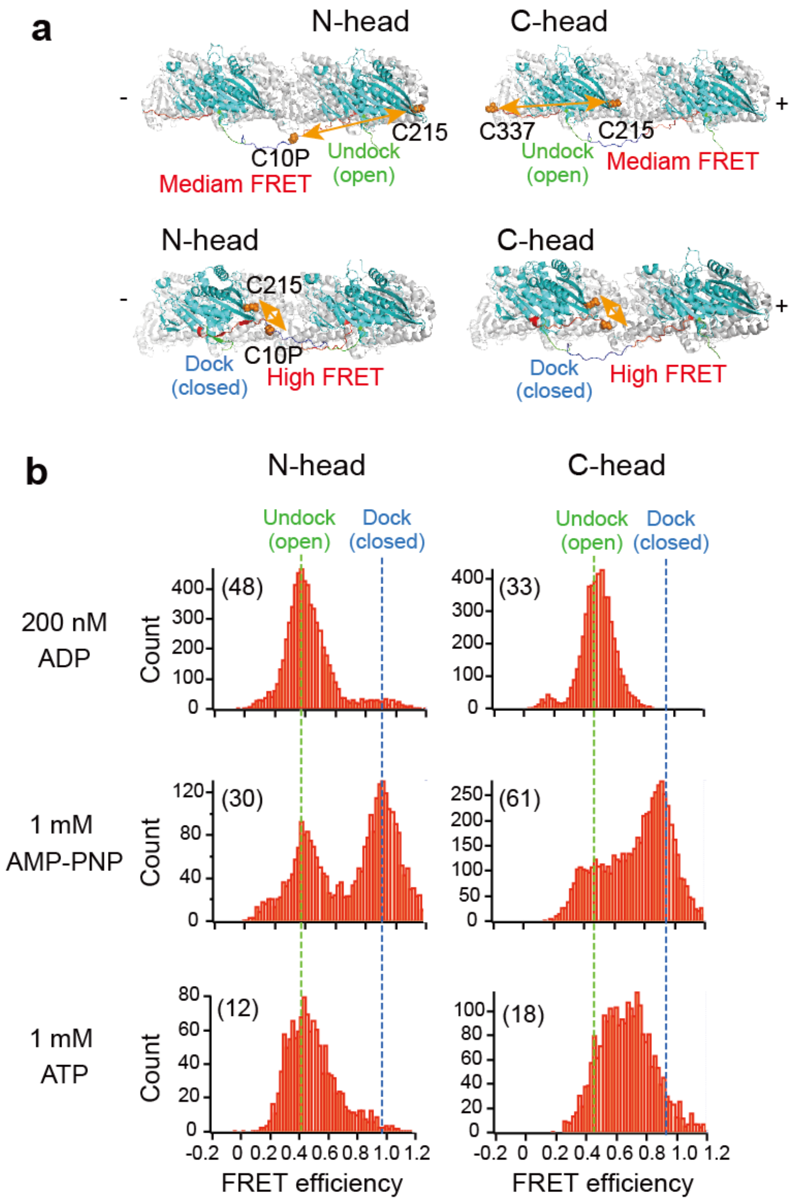


**Supplementary Figure 15 | SmFRET observations of neck linker conformational states of N- and C-heads of tandem kinesin.** (**a**) Diagrams illustrating the positions of cysteine residues for dye labeling. To detect the neck linker conformation of the N-head, E215 and the first proline residue of the 11 polyproline linker were replaced with cysteine (termed C10P; left). For the C-head's neck linker conformation, E215 was replaced with cysteine, and an additional cysteine was introduced after the neck linker at residue 337. (**b**) Histograms of FRET efficiencies (from each frame of images) of Cy3/Cy5-labeled tandem kinesin bound to the axonemes with 200 nM ADP, 1 mM AMP-PNP, and 1 mM ATP (typical traces are shown in Fig. 5b). Images were recorded either at 50 (for ADP and AMP-PNP conditions) or at 100 (for ATP) fps. Green and blue dotted lines illustrate putative undocked and docked states. Parentheses show the numbers of molecules analyzed. The smFRET sensor on C-head in 1 mM ATP showed a broad distribution of FRET efficiency between docked and undocked state, indicating that the neck linker transitions rapidly between these states (at speed comparable to or faster than the 10 ms temporal resolution of smFRET).

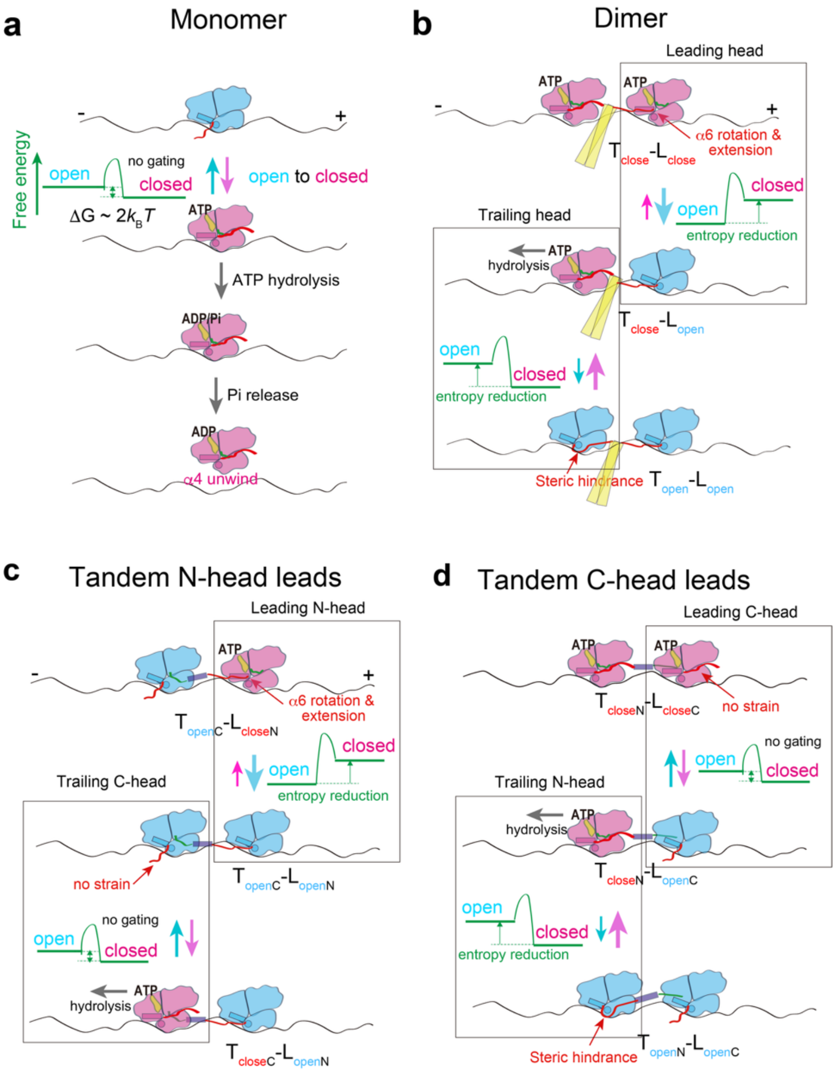


**Supplementary Figure 16 | Kinetic asymmetry in open-closed conformational transition explaining the preferential detachment of the trailing head from microtubule.** Schematic explaining the regulation of the microtubule-detachment of the monomer (**a**), leading and trailing heads of the dimer in two-head-bound state (**b**), leading N-head and trailing C-head (**c**), and leading C-head and trailing N-head (**d**) of tandem kinesin in two-head-bound states. Open and closed conformational states of kinesin head are indicated in cyan and magenta, respectively. Cover strand and neck liner are depicted in green and red. Free energy landscapes along the conformational coordinate are shown in green lines. Without neck linker constraint (panel **a**), the activation energies of the open-to-closed and closed-to-open conformational transitions differ only slightly (the free energy change upon neck linker docking is relatively small[5]), allowing the head to transition between open and closed states (without gating). In dimeric kinesin (panel **b**), when the leading head transitions from open to closed state, the α6 helix rotates and its C-terminus extends. This increases tension in the neck linker[1], destabilizing the closed state through entropy reduction (front-head gating). In contrast, when the trailing head transitions from closed to open state, steric hindrance just ahead of its base (C-terminus of α4 helix) increases tension in the neck linker[1], kinetically and thermodynamically stabilizing the closed state (rear-head gating). In tandem kinesin (panels **c**, **d**), the neck linker of C-head is not connected to its partner head and thus experiences no gating. Nevertheless, the N-head's front-head or rear-head gating alone is sufficient to maintain processive motility.
